# Haem toxicity in the aged spleen impairs T-cell immunity through iron deprivation

**DOI:** 10.1101/2024.05.05.592551

**Authors:** David Ezuz, Heba Ombashe, Lana watted, Orna Atar, Noga Ron-Harel

## Abstract

Ageing profoundly impacts T cell immunity, compromising vaccine responses, susceptibility to infections, and immunosurveillance. Mechanisms of T cell ageing involve cell-intrinsic alterations and interactions with other immune and stromal cells^1^. This study investigates how a tissue’s microenvironment influences T cell ageing trajectories, following our discovery of varying rates of T cell ageing within a single host. Spleen-derived lymphocytes exhibited functional decline compared to those from lymph nodes, with proteomic analysis revealing increased expression of haem detoxification and iron storage proteins in aged spleen-derived lymphocytes. Exposure to the aged spleen microenvironment or to haem induced multiple ageing phenotypes in young lymphocytes, characterized by reduced proliferation and upregulation of the ectonucleotidases CD39 and CD73. Mechanistically, we show that T cells survive the hostile microenvironment of the aged spleen by maintaining low labile iron pools to resist ferroptosis. Finally, vaccination responses in aged mice were enhanced by timed iron infusions. Our findings underscore a trade-off between T cell survival and function in the aged host. Recent studies show how Dysfunctional T cells induce premature ageing phenotypes in multiple solid organs of a young host^2,3^. Our findings highlight the bidirectional relationship between T cells and their ageing microenvironment. Understanding these mechanisms will inform strategies to enhance immune responses in the elderly.

## Introduction

T lymphocytes, the cellular arm of the adaptive immune response, play a pivotal role in host defence against foreign pathogens and contribute to maintaining tissue homeostasis. However, their functionality diminishes with age, leading to compromised immunity against infections, diminished response to vaccination, and an elevated susceptibility to autoinflammatory diseases and malignancies^1^. The mechanisms driving T cell dysfunction in ageing involve universal hallmarks of cellular ageing, such as mitochondrial dysfunction, loss of proteostasis, genetic alterations, and senescence^4^, together with T cell-specific hallmarks, including a reduction in T cell repertoire and a phenotypic shift towards less naive and more differentiated cells in elderly individuals^1^. Recent studies in mice show that the premature ageing of the T cell compartment, coupled with the subsequent failure of immunosurveillance, accelerates ageing phenotypes in multiple organs^2,3^. In this study, we explored the reciprocal nature of this interaction as we pursued our unexpected observation highlighting functional differences between naive T cells obtained from aged spleens to those obtained from peripheral lymph nodes within the same host. Specifically, T cells obtained from spleens demonstrated low viability, and reduced proliferation when stimulated ex vivo, compared to T cells derived from lymph nodes. In addition, T cells derived from the aged spleen showed overexpression of CD39, a cell surface ATPase previously associated with metabolic stress and reduced response to vaccination in T cells obtained from aged individuals^5^. Given that T cells could move in and out of circulation, migrating into various lymphoid organs, this observation raised intriguing questions: what factors within the aged spleen microenvironment were driving these inherent ageing phenotypes in its resident T cells, and through what mechanisms?

Secondary lymphoid organs (spleen and lymph nodes) play a vital role in maintaining the quiescent T cell pool and facilitating T cell-mediated immunity. With ageing, these organs undergo significant changes in tissue organization, cellularity, and function, which compromise their capacity to sustain T cell homeostasis and activation^6,7^. Structural changes include internal disorganization and fibrosis. In cell transfer experiments, young T cells ‘parked’ in an aged lymph node exhibit reduced levels of STAT5 phosphorylation, a homeostatic signal downstream of IL7 receptor, and lower survival compared to cells residing in young lymph nodes^6^. In response to infection or vaccination, an adult lymph node can expand 10-fold, whereas aged lymph nodes undergo modest expansion and never reach the same cellularity^8^. A similarly inferior response occurs even when young T cells are injected into an aged mouse^6^, suggesting dysfunction of the aged lymph node stromal cells. The spleen is organized in regions called the red pulp (RP) and white pulp (WP), separated by an interface called the marginal zone (MZ). The white pulp resembles the structure of a lymph node, containing T cell and B cell zones, while the red pulp primarily functions to filter blood and recycle iron from senescent red blood cells, a task predominantly carried out by red pulp macrophages^9^. Functional deterioration of red pulp macrophages with ageing leads to accumulation of senescent red blood cells, haem, and iron depositions, in aged spleens^10^. Excess haem and iron could promote oxidative stress and lipid peroxidation^11^. Thus, we hypothesized that such changes in the microenvironment of the aged spleen determine the ageing phenotypes of its resident T cells.

Using unbiased, whole-cell proteomics, we discovered heightened expression of stress- and inflammation-associated proteins in naïve T cells from aged spleens compared to young, including enzymes involved in haem catabolism and iron storage. Exposure to the aged spleen microenvironment or haem induced this distinctive protein signature and hinder proliferation in young T cells. We found that T cells survived in the aged spleen’s harsh conditions through resistance to ferroptosis, a cell death pathway driven by iron and reactive oxygen species (ROS); Both aged T cells and young T cells exposed to the aged spleen microenvironment exhibited enhanced survival upon ferroptosis induction compared to control young T cells. Mechanistically, ferroptosis resistance stemmed from depleted labile iron pools, leading to compromised proliferation. In vivo supplementation of aged mice with labile iron improved T cells response to vaccination.

## Results

### The aged spleen microenvironment induces T cell dysfunction

To investigate the impact of tissue microenvironment ageing on its resident T cells while controlling for differences arising from changes in the T cell population’s composition, we purified naïve CD4^+^ T cells (CD4^+^CD25^-^CD62L^+^CD44^lo^) from the spleen and lymph nodes (LNs) of aged (21-23 months old) C57Bl/6 mice and analysed their response to activation ex vivo (Fig. 1A). Significant functional differences were identified as aged T cells collected from spleens showed lower viability (Fig. 1B) and reduced proliferative capacity (Fig. 1C, 1D), compared to aged T cells residing in LNs. Expression levels of early activation markers (CD69, CD25) and the central co stimulatory molecule (CD28) were not substantially different (Extended data Fig. 1A-C). Importantly, naïve T cells derived from the spleen or LNs of young mice did not differ in their response to activation, demonstrating comparable viability and proliferation (Extended data Fig. 1D,E). CD39 is expressed on dysfunctional and exhausted T cells ^5,12^. Our analysis discovered a site-specific regulation of CD39 expression. CD39 levels were significantly higher in T cells derived from the spleen compared to LNs in both young and aged mice (Fig. 1E, 1F). The fact that this differential expression was more pronounced in aged T cells, suggested that signals inducing CD39 upregulation in the spleen microenvironment were exacerbated with ageing. To directly examine whether exposure to the aged spleen milieu, in vivo, was sufficient to induce functional defects, young T cells (tdTomato^+^) were transfused intravenously into young or aged recipients. 2-3 weeks following cell transfer, recipient mice were sacrificed, CD4^+^ T cells were harvested from their spleens and lymph nodes and analysed by flow cytometry (Figures 1G and Extended data Fig. 1F show experimental design and gating strategy, respectively). Young CD4^+^ T cells residing in aged spleens showed a significant increase in cell size (Fig. 1H), and elevated CD39 levels (Fig. 1I). Observed differences in CD39 expression maintained after activation (Extended data Fig. 1G-H). Moreover, in response to ex vivo stimulation, young CD4^+^tdTomato^+^ T cells parked in the aged spleens proliferated less than T cells purified from the LNs of the same host, or from a young host (Fig. 1J, 1K). Together, these results show that the microenvironment within the aged spleen promotes T cell dysfunction.

**Figure 1:**
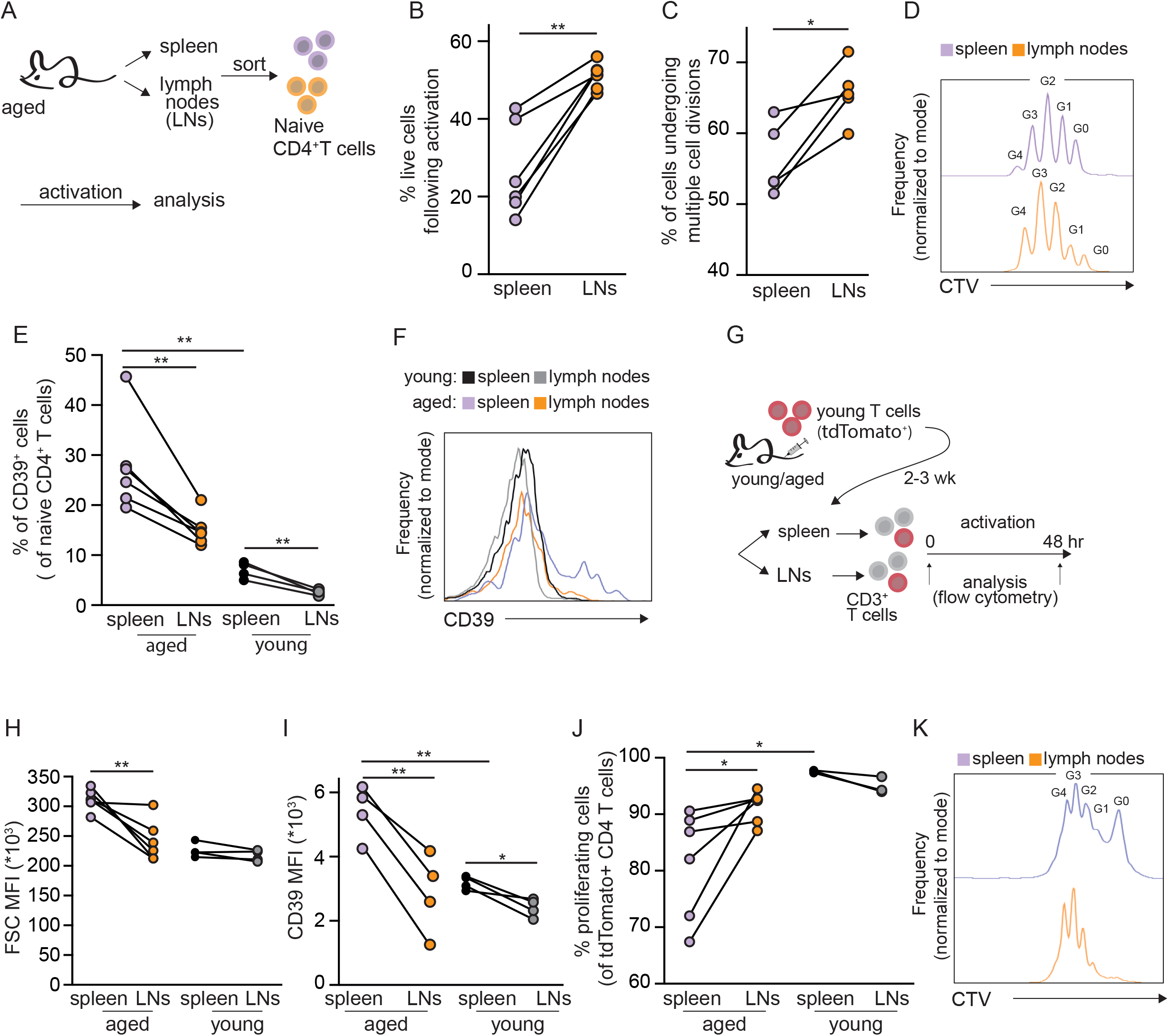
The aged spleen microenvironment induces T cell dysfunction. (A-D) (A) Scheme of experimental design. Naïve T cells (CD4+CD25-CD62L+CD44lo) were purified from the spleen or peripheral lymph nodes (LNs) of aged mice (20-22 months old). The cells were kept in two separate pools and stimulated ex vivo for 48 hr using plate bound anti-CD3/anti-CD28. (B) Cell viability, analyzed by flow cytometry. (C) Quantitation (D) a representative FACS histogram demonstrating cell proliferation, detected by dilution of CellTrace Violet. (E) Quantification, and (F) a representative FACS histogram showing CD39 expression on T cells isolated from the spleen and LNs of young (8 weeks old) and aged mice, analyzed by flow cytometry. (G) Scheme of experimental design for T cell adoptive transfer. Young T cells derived from TdTomato+ transgenic mice were transfused into young or aged C57Bl/6 wild type recipients. After 2-3 weeks, recipient mice were sacrificed, and CD4+ T cells purified from the spleen and LNs for analysis of (H) cell size and (I) CD39 expression by flow cytometry. (J, K) A portion of the cells were loaded with CellTrace violet, and stimulation for 48 hr, to assess proliferation. Each data point represents an individual mouse. (*p<0.05, **p<0.01; paired student’s t test when comparing cells derived from spleen and LNs of the same mouse, and unpaired student’s t test when comparing values across age groups). Each panel shows representative data of at least 2 independent experiments.

### T cells in the aged spleen are expressing high levels of proteins closely associated with stress and inflammation

To identify T cells’ cellular response to the aged spleen microenvironment, we performed whole-cell, label-free proteomics analysis of pure naive CD4^+^ T cells collected from the spleens of young and aged mice. Cells were immediately processed or stimulated ex vivo for 24 hr prior to protein extraction, peptide degradation and analysis by LC-MS/MS (Fig. 2A). The dynamic changes of over 3800 proteins were determined (Extended data Table 1). According to principal component analysis (PCA), the largest changes in protein composition were induced by activation (PCA1 represents 54% of the variance and separates between 0 and 24 hr). Age accounted for 17.2% of the variance (PCA2, separating young and aged T cells; Fig. 2B). 692 proteins were differentially expressed between young and aged T cells post-activation. 84% of those were higher in young T cells (Fig. 2C). In agreement with previous studies showing metabolic defects in aged T cells ^13,14^, pathway enrichment analysis highlighted multiple metabolic pathways that were significantly over-represented in young vs aged T cells, including: one-carbon metabolism, purine and pyrimidine metabolism, and amino acids metabolism (Fig. 2D). We further identified enrichment of proteins involved in DNA replication and protein translation (Fig. 2D), in agreement with the deficient growth and proliferation of aged T cells (Fig. 1). Despite their naïve identity and the strict sorting parameters of naïve T cells, 330 proteins were differentially expressed between naïve young and aged T cells (Fig. 2E). 87% of them were overrepresented in aged T cells and were enriched with proteins closely associated with activation, proinflammation, and stress responses (Fig. 2F). Most proteins over-expressed in aged naïve T cells are normally induced by activation in young T cells, as demonstrated by the overlap between the two protein groups (Fig. 2G). Thus, despite expressing surface markers characteristic of naïve T cells, the proteome of aged T cells suggest engagement in multiple stress responses. Many of these proteins remained higher in aged T cell even after activation (Extended data Fig. 2A), suggestive of enduring and inherent changes in the aged T cells. To identify specific pathways induced in aged naive T cells compared to young, we performed pathway enrichment analysis of the top 100 differentially expressed proteins. This analysis highlighted proteins associated with haem metabolism and degradation (Fig. 2H). Haem is catabolized by haem oxygenase 1 (HMOX1 or HO-1) to generate carbon monoxide (CO), biliverdin, and labile iron. Excess iron is stored in cells within the cavity of ferritin, a globular hollow protein composed of 24 subunits of two types: ferritin H (FTH1) and ferritin L (FTL). Biliverdin is further reduced to bilirubin by biliverdin reductase (BLVRB) (Fig. 2I). All these proteins were significantly elevated in aged naïve CD4^+^ T cells compared to young (Fig. 2H). Overexpression of HO-1 (Fig. 2J, 2K) and ferritin (Fig. 2L, 2M) was further verified by flow cytometry. We postulated that all T cells residing in the aged spleen were similarly exposed to any age-related signals. Indeed, qPCR analysis comparing bulk CD3^+^ T cells from the spleens of young and aged mice was performed, indicating increased expression of *Ho-1* (Extended data Fig. 2B), the two genes encoding for biliverdin reductase isozymes: *Blvra* and *Blvrb* (Extended data Fig. 2C-D, respectively), and the two ferritin subunits: *Ftl* and *Fth1* (Extended data Fig. 2E,F) in aged T cells. Haem makes an interesting candidate when looking for a deleterious signal that is specifically affecting T cells in the aged spleen and not LNs, as it is directly connected to the spleen being a site of senescent red blood cells (RBC) removal and iron recycling by red pulp macrophages ^9^.

**Figure 2:**
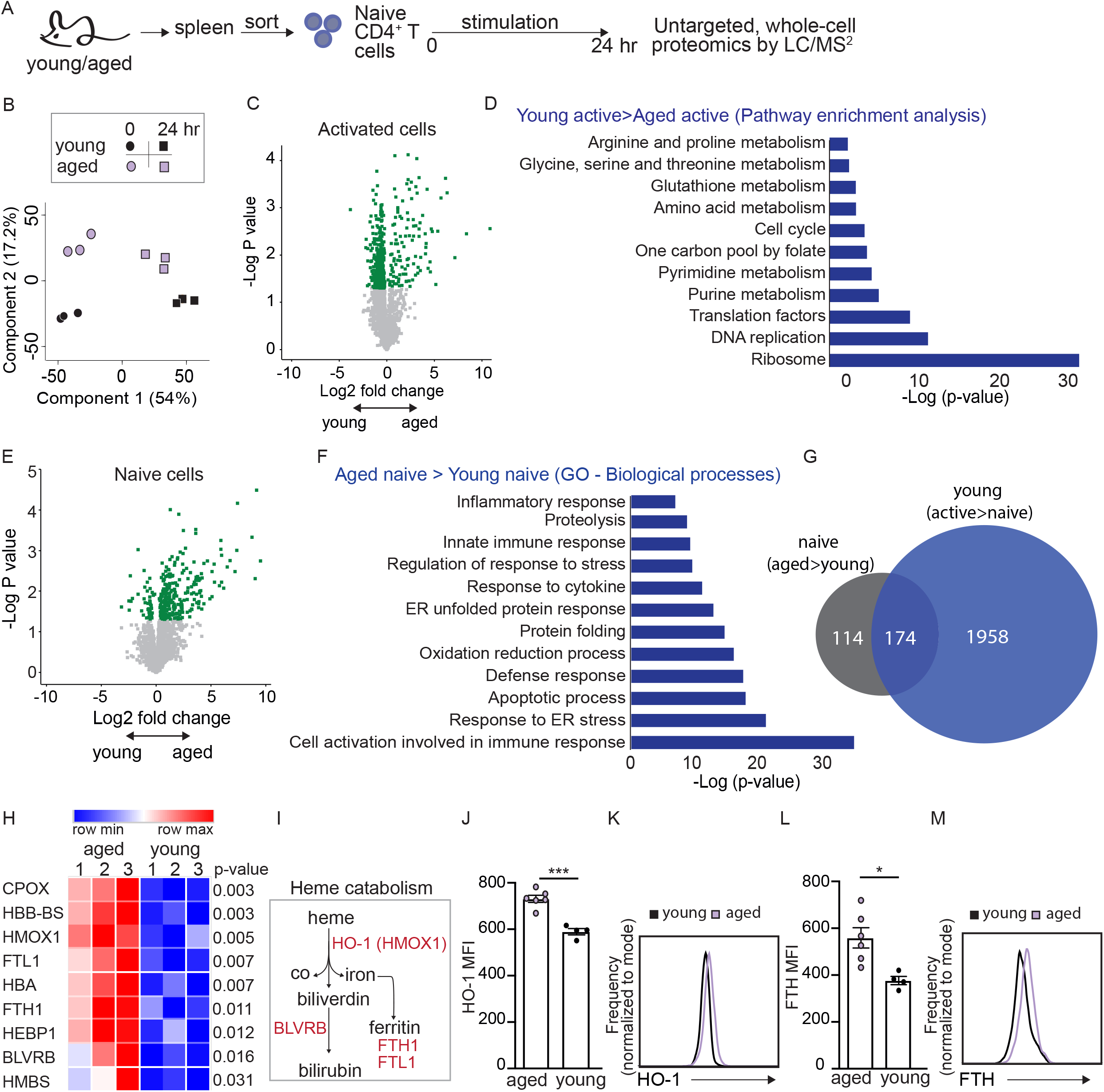
T cells in the aged spleen are expressing high levels of proteins closely associated with stress and inflammation. (A) Experimental scheme. Naïve CD4+ T cells were sorted from the spleen of young (n=3) and aged mice (n=3 pools of 2 mice) and were either immediately frozen or stimulated ex vivo using plate bound anti-CD3/anti-CD28 antibodies prior to protein extraction and digestion. The peptide pool from each sample was analyzed by LC-MS/MS. (B) Principal component analysis. (C) Volcano plot showing differences in protein levels between activated young and aged T cells. Green signifies statistical significance. (D) Pathways enriched among proteins significantly overrepresented in young vs aged T cells following activation. (E) Volcano plot showing differences in protein levels between naive young and aged T cells. Green signifies statistical significance. (F) Pathways enriched among proteins significantly overrepresented in aged vs. young naïve T cells. (G) Venn Diagram showing overlap between proteins overexpressed in aged naïve T cells and those elevated with young T cell activation. (H) Heatmap summarizes proteins associated with heme metabolism and detoxification, elevated in aged naïve T cells compared to young. (I) Key proteins involved in heme catabolism. (J) Analysis of HO-1 Mean fluorescent intensity (MFI; Geometric Mean). (K) a representative plot. (L) Analysis of FTH Mean fluorescent intensity, and (M) a representative plot. Bar graphs represent mean ± SEM. Data points represent single mice (*P<0.05, ***P<0.001; unpaired student’s t-test).

### T cells in the aged spleen microenvironment are exposed to toxic haem and iron depositions

Following our observations of site-specific T cell dysfunction (Fig. 1) and the identification of the intracellular response to haem (Fig. 2) we investigated whether aged T cells were indeed exposed to high haem concentrations in vivo by quantifying haem in the splenic microenvironment and its resident T cells. We detected elevated levels of haem in CD3^+^ T cells isolated from aged spleens (intracellular haem; Fig. 3A), and in the interstitial fluid enriched fraction derived from aged spleens compared to young (extracellular haem; Fig. 3B). The interstitial fluid enriched fraction of aged spleens further contained higher levels of secreted unconjugated bilirubin, a product of haem degradation (Fig. 3C). To directly test whether exposure to an aged spleen microenvironment in vivo was sufficient to induce the protein signature of aged T cells (Fig. 2), young T cells derived from tdTomato transgenic mice were transferred into young and aged recipients. After 2-3 weeks, the mice were sacrificed and CD3^+^ T cells were purified and analysed by flow cytometry, gating on tdTomato^+^ T cells (Fig. 3D). Strikingly, young T cells residing in the aged spleen but not lymph nodes significantly upregulated the FTH1 subunit of ferritin (Fig. 3E, 3F) and HO-1 (Fig. 3G, 3H). With ageing, the spleen infrastructure becomes compromised, and the borders between the red pulp and white pulp become less clear (Fig. 3L, and ^15^). The white pulp is surrounded by marginal zone macrophages, that can be identified by expression of CD169 (also known as Sialoadhesin or Siglec1). Immunohistochemical analysis of CD169 expression further affirmed the observed loss of tissue organization, and compromised separation between the two regions in aged spleens (Fig. 3M). Our H&E staining identified brown depositions in aged spleen tissues (Fig. 3L). Previous studies determined those were iron-rich aggregates, indicative of high iron load in the aged spleen, due to inefficient recycling of senescent red blood cells in the aged tissue ^10^. Thus, we applied the Prussian Blue method to detect iron depositions in the spleen. As previously reported, the signal was higher in aged tissues, and could be detected also inside white pulp areas (Figure 3N). Together, these data suggest that failure of senescent red blood cell recycling in the aged spleen, together with loss of tissue organization expose T cells to a potentially toxic microenvironment, enriched with haem and iron depositions. In agreement, addition of the interstitial fluid enriched fraction of an aged spleen (SE) to young T cells in culture promoted cell death, in a dose-dependent manner. The cells survived significantly better in SE derived from young mice (Extended data Fig. 3A). Addition of Desferoxamine (DFO), an iron chelator, partially rescued cell viability, in cells treated with high concentrations of aged SE (1:4 dilution in growth media; Fig. 30, 3P).

**Figure 3:**
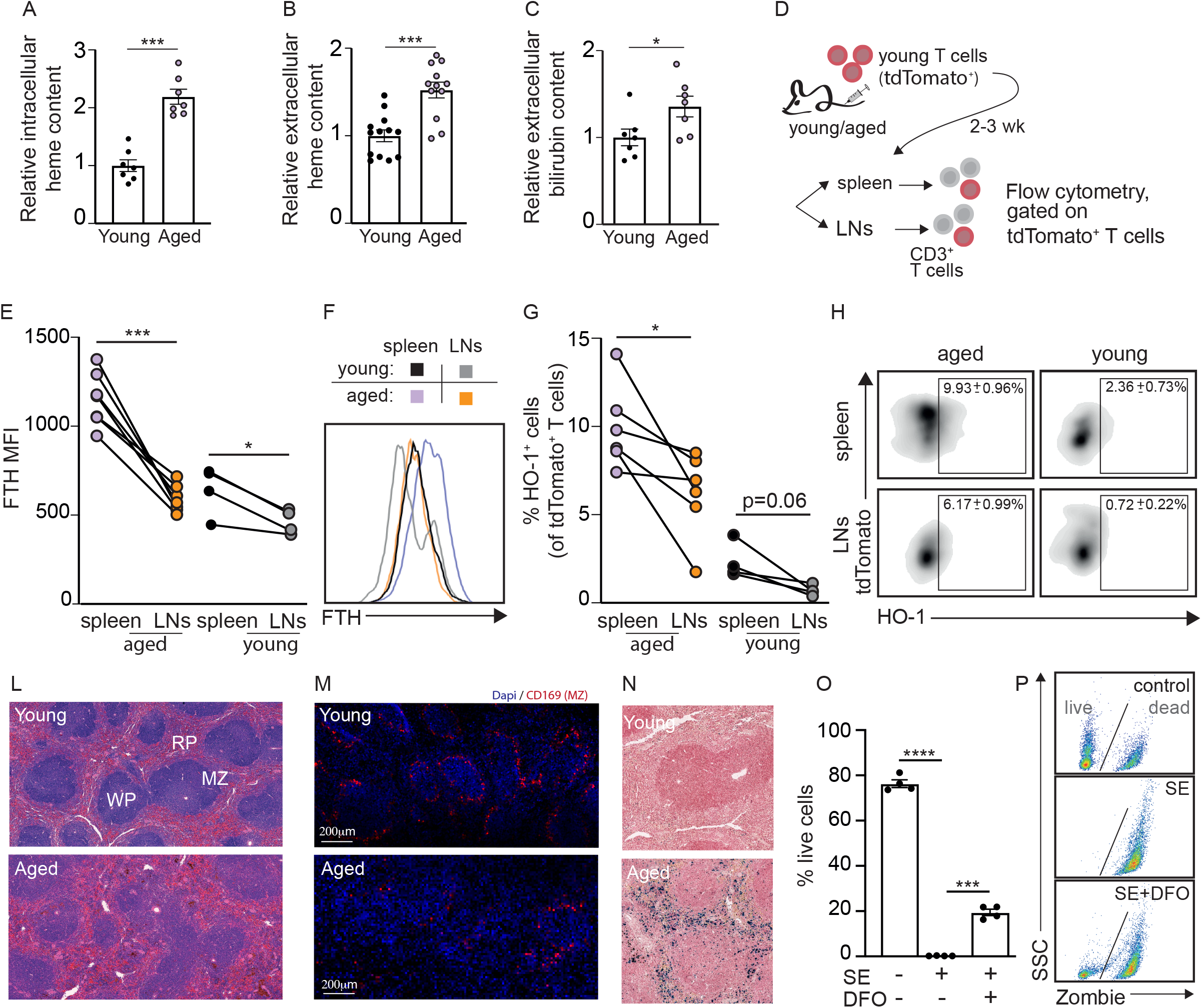
T cells residing in the aged spleen are exposed to toxic heme and iron depositions. Spleens were excised from young and aged mice. (A) Heme content was calorimetrically quantified intracellularly, in isolated CD3+ T cells. The interstitial-fluid enriched fraction was used to quantify (B) heme and (C) bilirubin levels in the tissue microenvironment. (D) Scheme of experimental design. Young T cells derived from transgenic mice ubiquitously expressing TdTomato were transferred into young or aged C57Bl/6 wild type recipients. After 2-3 weeks, recipient mice were sacrificed. CD3+ T cells were purified from the spleen and lymph nodes, and stimulated for 48 hr ex vivo, prior to analysis by flow cytometry to quantify (E,F) FTH and (G,H) HO-1 expression levels by flow cytometry. (L) H&E processed paraffin sections of spleens derived from aged and young mice (RP:red pulp; WP:white pulp; MZ: marginal zone). (M) representative images of frozen spleen sections stained with anti-CD169 to mark MZ macrophages. (N) Spleen paraffin sections from aged and young mice, processed by Prussian Blue method to detect ferric iron depositions in the spleen. (O) Young T cells were treated with aged spleen’s interstitial-fluid enriched fraction (SE) in high concentrations ± desferoxamine (DFO), an iron chelator. Cell viability was quantified by flow cytometry, using Zombie Aqua staining. (P) Representative flow plots. Bar graphs represent mean ± SEM. Data points represent single mice (*P<0.05, ***P<0.001, ****P<0.0001; paired student’s t test when comparing cells derived from spleen and LNs of the same mouse, and unpaired student’s t test when comparing values across age groups). Each panel shows representative data of at least 2 independent experiments. Panels A-C shows pooled data from 3 independent experiments.

### Haem drives ageing-like phenotypes in young T cells

To examine whether haem itself could drive T cell dysfunction, CD3^+^ T cells were purified from spleens of young mice and stimulated using plate bound anti-CD3/anti-CD28 antibodies, in the presence of haem, with or without bovine serum albumin (BSA), which binds and sequesters haem and its by-products^16^. Cell viability was reduced in cells exposed to haem at high concentration, and partially rescued by addition of BSA (Fig. 4A,B). Moreover, haem inhibited T cell proliferation and its effect was almost completely reversed by addition of BSA (Fig. 4C,D). BSA further rescued proliferation in T cells incubated with high concentrations of SE (1:4 dilution in culture media; Extended data Fig. 4A,B). Haem degradation products regulate T cell functions^17,18^. To test whether exposure to haem itself could impair T cells activation, T cells were treated with haem in combination with tin-protoporphyrin (SnPP), an HO-1 inhibitor. SnPP caused proliferation arrest even under lower concentrations of haem (50μM; Fig. 4E). Moreover, exposure to haem was sufficient to induce CD39 levels in young T cells (Fig. 4F, 4G). Like proliferation arrest, CD39 expression was directly induced by haem, as addition of SnPP caused an even greater boost in CD39 levels (Fig. 4H, 4I). Thus, exposure of young T cells to haem caused multiple phenotypes seen in T cells residing in an aged spleen, in vivo. Haem and iron accumulate in the aged spleen due to inefficient recycling of senescent red blood cells (RBC) and increased probability for red blood cell lysis within the tissue^10^. This could lead to an increase in extracellular ATP accumulation, a potent proinflammatory, danger signal that regulates T cell function^19,20^. CD39, together with an additional ectonucleotidase, CD73, convert ATP to ADP, AMP and adenosine^21^. Like CD39, CD73 levels were significantly upregulated in aged, spleen-derived T cells compared to young (Extended data Fig. 4C). Moreover, young TdTomato^+^ T cells that were transferred into an aged host overexpressed CD73 compared to TdTomato^+^ T cells transferred into a young host (Extended data Fig. 4D). In agreement, LC-MS analysis of the interstitial-fluid enriched fraction (SE) derived from young and aged spleens, showed an increase in ATP and adenosine levels in aged SE (Extended data Fig. 4E). Accumulation of extracellular adenosine suppresses T cell activation^22,23^, offering a potential mechanism for T cell dysfunction within the aged spleen microenvironment. However, aged T cells exhibited defects in proliferation, even when removed from the aged spleen. This raised the question: What lasting cellular impacts stem from exposure to the aged spleen microenvironment?

**Figure 4:**
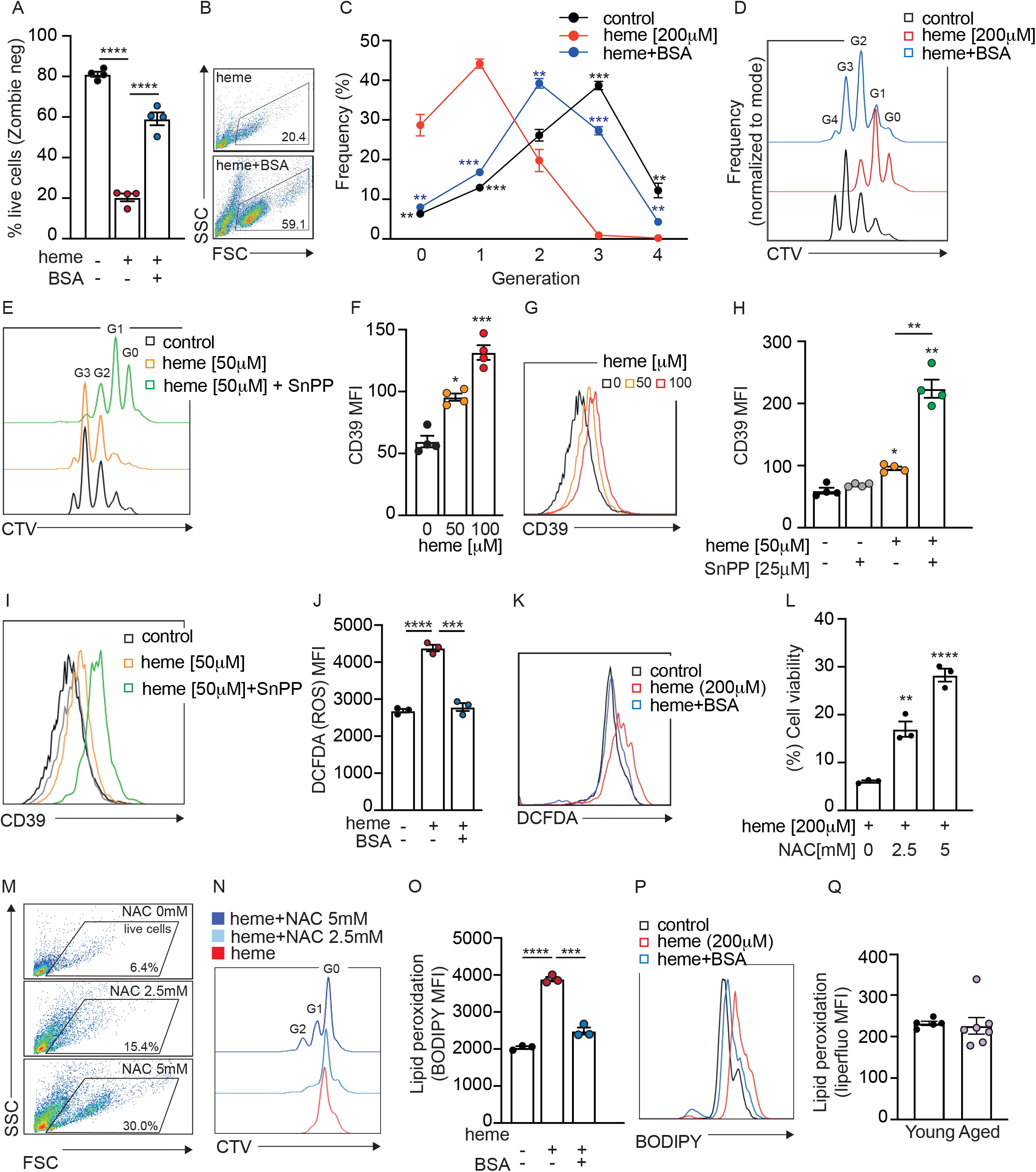
Heme promotes aging-like phenotypes in young T cells. Young T cells were loaded with CellTrace Violet (CTV), cultured in media supplemented with heme ± bovine serum albumin (BSA) for 48 hr, and analyzed by flow cytometry to assess (A,B) cell viability and (C, D) proliferation. Gn indicates number of cell divisions. Statistics was calculated on cells treated with heme compared to heme+BSA or control cells. (E) A representative plot showing proliferation of cells treated with heme ± tin protoporphyrin IX (SnPP), an HO-1 inhibitor. (F,G) Analysis of CD39 expression levels in young T cells treated with heme. (H, I) Analysis of CD39 expression levels in cells treated with heme ± SnPP. (J,K) dichlorodihydrofluorescein diacetate (DCFDA) was used to quantify ROS in young T cells cultured in media supplemented with heme ± BSA for 48 hr. Young T cells were cultured with heme± NAC for 48 hr and analyzed by flow cytometry to assess (L, M) viability and (N) proliferation. (O,P) Analysis of BODIPY fluorescence intensity, as an indication of lipid peroxidation in young T cells treated with heme ± BSA. (Q) Analysis of lipid peroxidation using the liperfluo reagent in young and aged T cells. Bar graphs represent mean ± SEM. Data points represent individual mice (*P<0.05, **P<0.01, ***P<0.001, ****P<0.0001; unpaired student’s t-test, comparing each treatment to control cells, unless otherwise indicated). Each panel shows representative data of at least 2 independent experiments.

Haem is a potent inducer of reactive oxygen species (ROS) ^24^. Indeed, T cells exposed to haem showed an increase in cellular ROS, which was completely abolished by BSA (Fig. 4J,K). Addition of the antioxidant N-acetyl cysteine (NAC) improved cell viability (Fig. 4L,M) and proliferation (Fig. 4N), suggesting that haem-induced ageing phenotypes in young T cells were partly mediated by ROS. Similarly, NAC improved cell viability and proliferation of T cells cultured with SE (Extended data Fig. F-H).

Excessive ROS promotes lipid peroxidation and cell death by ferroptosis^11^. In agreement, lipid peroxidation was elevated in T cells exposed to haem, and rescued by BSA (Fig, 4O,P). Taken together, these results suggested that haem accumulation in the aged spleen microenvironment induced oxidative stress in T cells and could lead to cell death by ferroptosis. Yet, to our surprise, neither BSA nor NAC improved survival or proliferation in aged T cells (Extended data Fig. 4I, J, respectively). Moreover, quantification of lipid peroxidation in aged T cells compared to young showed no apparent difference (Fig. 4Q). Thus, components accumulating in the aged spleen microenvironment and specifically haem, induce ROS, lipid peroxidation, cell death and proliferation arrest in young T cells. Yet, aged T cells residing in the aged spleen seem to be resistant to lipid peroxidation and they do persist in vivo. We thus hypothesized that aged T cells developed mechanisms that allowed them to survive the hostile microenvironment of the aged spleen.

### T cells residing in the aged spleen develop resistance to ferroptosis

To survive in an aged spleen, resident T cells would need to develop resistance to ferroptosis. To test this hypothesis, CD3^+^ T cells were isolated from spleens of young and aged mice and treated with RSL3 ((1S,3R)-RSL3), a compound that induces ferroptosis by inhibition of glutathione peroxidase 4. Exposure to RSL3 promoted cell death in young T cells, and was counteracted by Ferrostatin or Liproxstatin, two commercially available inhibitors of ferroptosis (Extended data Fig. 5A). Strikingly, aged T cells survived this treatment significantly better than young T cells, even at high RSL3 concentrations (Fig. 5A). Furthermore, T cells derived from aged spleens survived RSL3 treatment significantly better than T cells derive from LNs of the same donors (Fig. 5B) and were the population most resistant to lipid peroxidation following RSL3 treatment (Fig. 5C).

**Figure 5:**
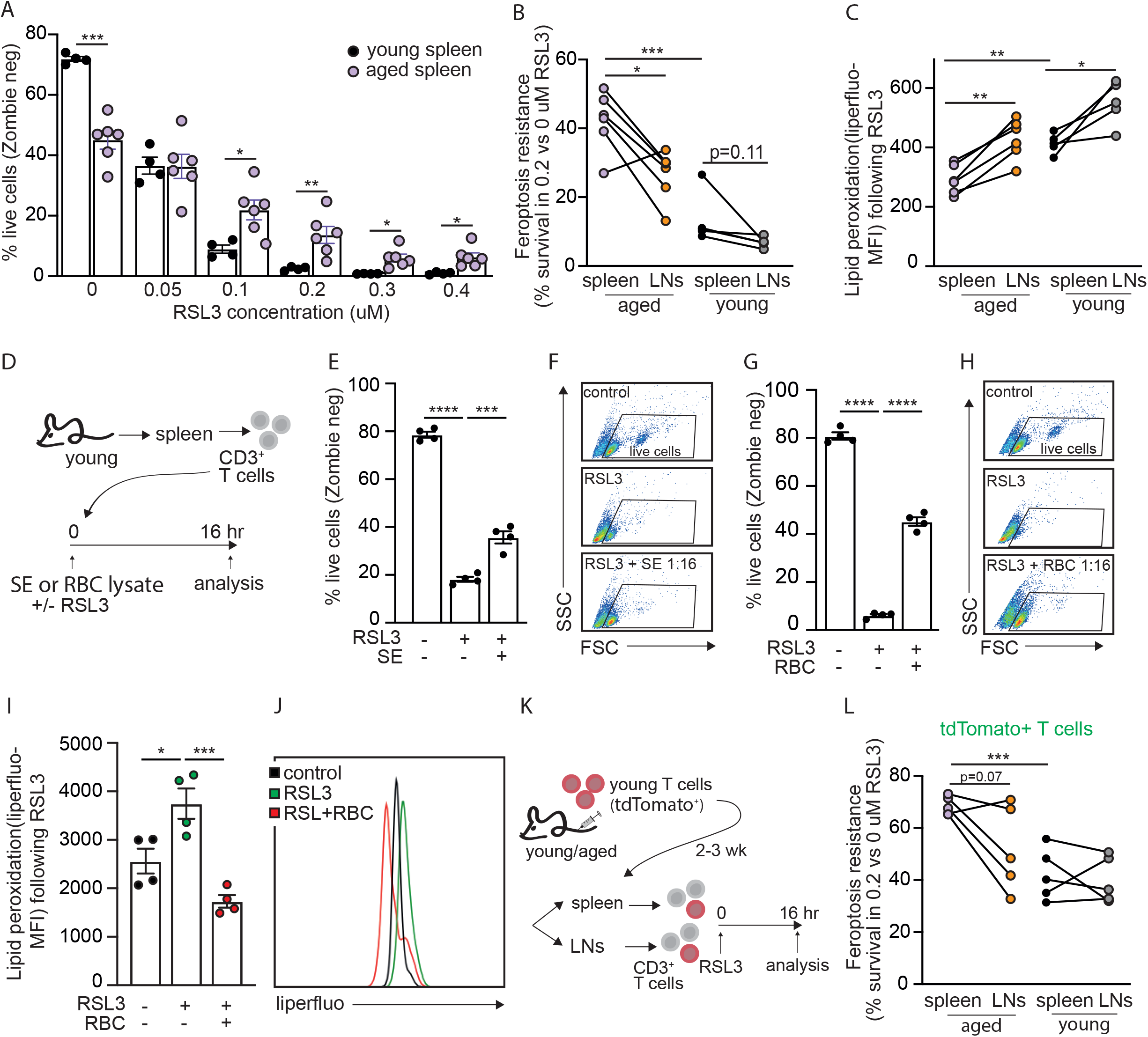
T cells residing in the aged spleen develop resistance to ferroptosis. T cells were purified from the spleens of young or aged mice and treated with increasing concentrations of RSL3. (A) Cell survival was quantified using the Zombie Aqua reagent, by flow cytometry. (B) Analysis of ferroptosis resistance (calculated as in A), and (C) lipid peroxidation (assessed using the Liperfluo dye) in young and aged T cells derived from spleens and lymph nodes. (D) Experimental scheme: Young CD3+ T cells were cultured ex vivo in the presence of RSL3 ± an aged spleen’s interstitial-fluid enriched fraction (SE) or red blood cell (RBC) lysate for 16 hr, following analysis by flow cytometry. (E-H) Analysis of cell viability under the different conditions. (I) quantitation and (J) a representative plot showing lipid peroxidation in response to RSL3 treatment ± RBC lysate. (K) Schematic of experimental design. Adoptive cell transfer was performed as described in Figure 1G. Purified T cells were cultured for 16 hr with RSL3 prior to analysis by flow cytometry to assess (L) ferroptosis resistance of transferred TdTomato+ T cells, analyzed as the relative viability in 0.2 vs 0 M RSL3. Bar graphs represent mean ± SEM. Data points represent single mice (*P<0.05, **P<0.01, ***P<0.001, ****P<0.0001; paired student’s t test when comparing cells derived from spleen and LNs of the same mouse, and unpaired student’s t test when comparing values across age groups or treatments). Each panel shows representative data of at least 2 independent experiments.

To examine if young T cells exposed to the aged spleen microenvironment developed resistance to ferroptosis, we first examined how exposure to the interstitial-fluid enriched fraction of the aged spleen (SE) ex vivo affected young T cells response to RSL3 (Fig. 5D). Strikingly, addition of SE to cell culture increased young T cells resistant to RSL3-induced ferroptosis by nearly two folds (from 18.2% to 35.7% survival: Fig. 5E, 5F). The outcomes of exposing young T cells to RBC lysate were like those of SE: We found a dose-dependent reduction in viability of T cells exposed to RBC lysate (Extended data Fig. 5B). Cell that survived this treatment were resistant to ferroptosis (Fig. 5G, 5H and S5C) and lipid peroxidation (Fig. 5I, 5J), suggesting that red blood cells could be the source of the signal promoting ferroptosis resistance.

Finally, to directly test whether exposure to the spleen microenvironment in vivo was sufficient to induce ferroptosis resistance in its resident T cells, we performed adoptive cell transfer of TdTomato^+^ T cells from young donors into young or aged recipients. 2-3 weeks following cell transfer, recipient mice were sacrificed, CD3^+^ T cells were collected from the spleen and lymph nodes, and the response of TdTomato^+^ T cells to RSL3 was analysed (Fig. 5K). Strikingly, the T cells most resistant to ferroptosis were those derived from the aged spleen. In most aged mice (4 out of 5), T cells derived from the spleen were more resistant to ferroptosis compared to T cells derived from the LNs (Fig. 5L). Together, these results show that T cells residing in the aged spleen adapt to their microenvironment by becoming resistant to ferroptosis.

### Aged T cells fail to induce labile iron pools for activation

We hypothesized that ferroptosis resistance was accomplished by maintaining low cellular iron levels. To measure changes in labile iron in T cells following activation, we employed FerroOrange, a fluorescent probe that interacts with cellular ferrous ions (Fe^+2^). In young T cells, labile iron pools increased as early as 9 hr post-activation and continued rising until 72 hrs (Fig. 6A). In agreement with our hypothesis, aged T cells failed to properly increase cellular iron following activation (Fig. 6B, 6C). Notably, this age-related drop in labile iron pools was more pronounced in CD4^+^-compared to CD8^+^-T cells, although both cell populations had lower iron compared to young cells. FerroOrange MFI dropped 30% in aged CD4^+^ T cells compared to young, and 15% in aged CD8^+^ T cells (Fig. 6D). To avoid iron overload, an increase in cellular labile iron is often accompanied by upregulation of ferroportin, the only known iron exporter ^25^. While baseline levels of ferroportin were comparable between aged and young T cells (Extended data Fig. 6A), upon activation, ferroportin levels were lower in aged T cells, consistent with their diminished induction in labile iron pools (Fig. 6E). A major pathway for increasing cellular iron is the uptake of transferrin-bound iron, mediated by the transferrin receptor (TFR1), also known as CD71. Young T cells upregulated CD71 immediately upon activation (Extended data Fig. 6B, and ^26^). In agreement with observed differences in labile iron, CD8^+^ T cells expressed higher levels of CD71 compared to CD4^+^ T cells, and only CD4^+^ T cells downregulated CD71 with ageing (Fig. 6F).

**Figure 6:**
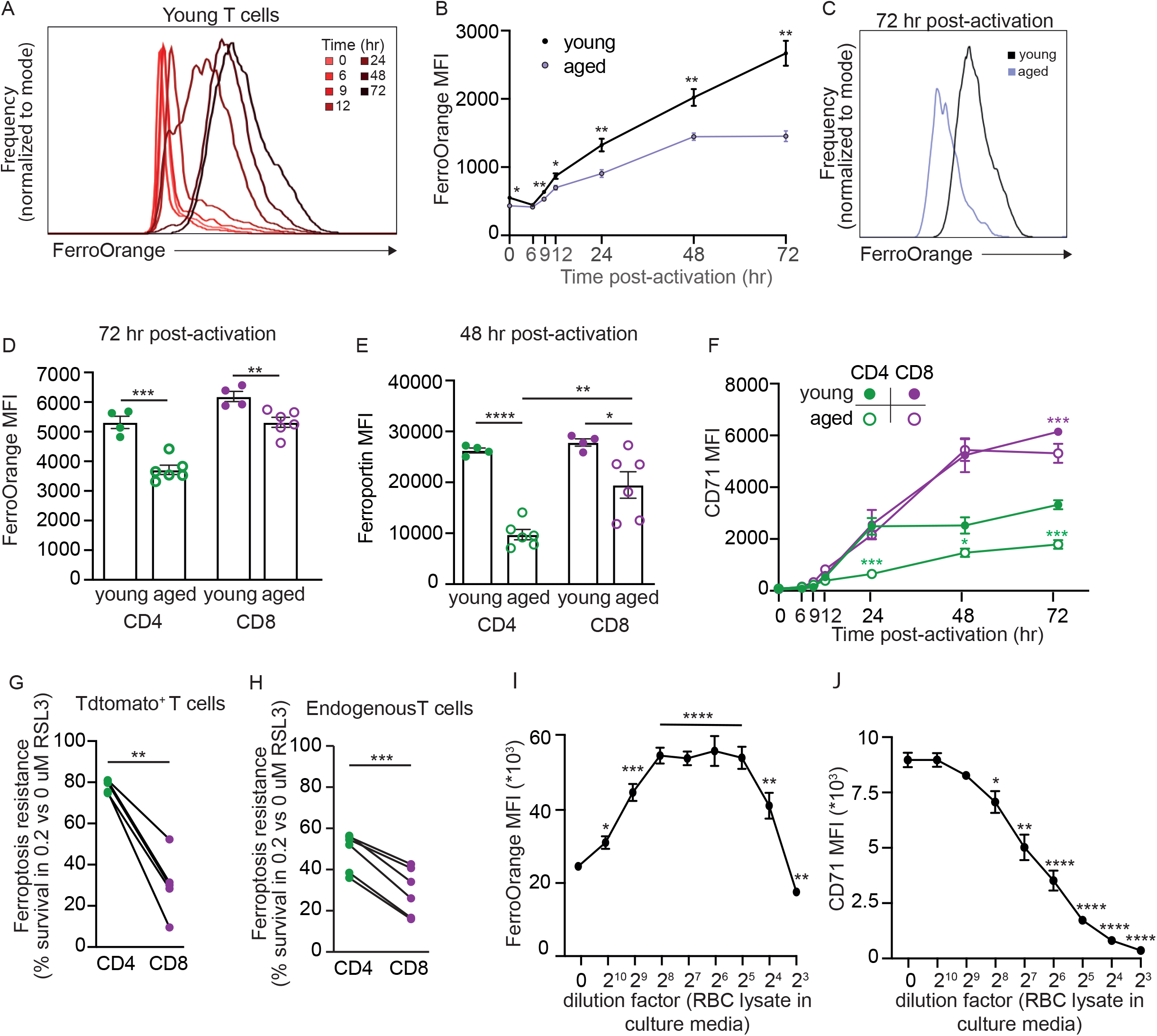
Aged T cells are iron deficient. (A) Young T cells were stimulated ex vivo using plate bound anti-CD3/anti-CD28. Cells were collected at different times post-activation) and loaded with Ferro-Orange, for detection of labile iron. (B) Kinetic changes in labile iron levels in young and aged T cells following activation. (C) a representative plot depicting differences in Ferro-Orange intensity at 72 hr post-activation. (D) Activation-induced differences in Ferro-Orange intensity in young vs aged CD4+ and aged CD8+ T cells. (E) Activation-induced differences in Ferroportin expression in young and aged CD4+ vs CD8+ T cells. (F) Analysis of CD71 (transferrin receptor 1) expression levels in young and aged T cells, following activation. (G, H) Re-analysis of experiment described in Fig. 5G, showing differences in ferroptosis resistance between Transferred (TdTomato+; G) and endogenous (TdTomato- ; H) CD4+ and CD8+ T cells. (I, J) Young T cells were activated ex vivo in media containing increasing doses of red blood cell (RBC) lysate. Labile iron (Ferro Orange; I) and CD71 expression (J) were analyzed by flow cytometry. Line graphs represent mean ± SEM. Data points (D,E,G,H) represent single mice . (*P<0.05, **P<0.01, ***P<0.001, ****P<0.0001; B,D,E,F: unpaired student’s t-test; G-H: paired student’s t-test; I-J: unpaired student’s t-test, comparing each condition to control cells).

To further connect low cellular iron to ferroptosis resistance, we reanalysed the adoptive transfer experiment (injecting young TdTomato+ T cells into young and aged hosts) shown in figure 5G, to assess ferroptosis resistance in specific T cell populations derived from the aged spleen. Strikingly, CD4^+^ T cells were more resistant to RSL3-induced ferroptosis compared to CD8^+^ T cells. This was seen in both TdTomato^+^ T cells (young T cells residing in the aged spleen; Fig. 6G) and TdTomato-T cells (aged endogenous T cells; Fig. 6H). To further establish the causal relation between ferroptosis resistance, iron uptake and cellular labile iron pools, young T cells were activated in media supplemented with increasing concentrations of RBC lysate. Activation induced iron uptake, and cells cultured in low levels of RBC lysate increased cellular iron pools even more, in agreement with RBC lysate being a rich source of iron. However, as the concentration of added RBC lysate increased, labile iron pool was reduced (Fig. 6I), together with a prominent downregulation of CD71 (Fig. 6J). Our findings highlight depletion of labile iron pools as a mechanism to resist ferroptosis in the aged spleen microenvironment.

### Iron supplementation rescued vaccination responses in aged T cells

Iron is essential for DNA synthesis and replication. Hence, we hypothesized that iron deficiency impairs proliferation in aged cells (as demonstrated in Fig. 1). To examine our hypothesis, we first tested whether labile iron pools in aged T cells could be replenished by supplementing them with ferric ammonium citrate (FAC), an iron derivative that bypasses transferrin receptor. Aged T cells were activated ex vivo in the presence of FAC. Analysis of cellular iron levels using FerroOrange verified that FAC increased labile iron pools in treated aged T cells (Fig. 7A, 7B). Additional indications of increased iron content were the downregulation of CD71 (Extended data Fig. 7A) and upregulation of ferroportin (Extended data Fig. 7B) in T cells supplemented with FAC. To examine whether iron supplementation could rescue proliferation in aged T cells, cells were stimulated ex vivo in the presence of FAC or iron saturated transferrin (holo-transferrin). Both treatments led to a significant improvement in proliferative capacity compared to non-treated aged T cells (Fig. 7C, 7D). In young T cells, iron supplementation did not affect proliferation (Fig. 7E), suggesting that iron was not a limiting factor in these cells. In agreement, iron supplementation increased EDU incorporation in activated aged but not young T cells (Extended data Fig. 7C). Young T cells ‘parking’ in an aged spleen for 2-3 weeks showed ferroptosis resistance (Fig. 5H) and impaired proliferation (Fig. 1H). Similar to our observations regarding endogenous young and aged T cells, iron supplement did not affect proliferation of young T cells residing in young spleens, but it significantly improved proliferation of young T cells residing in aged spleens (Extended data Fig. 7D-G).

**Figure 7:**
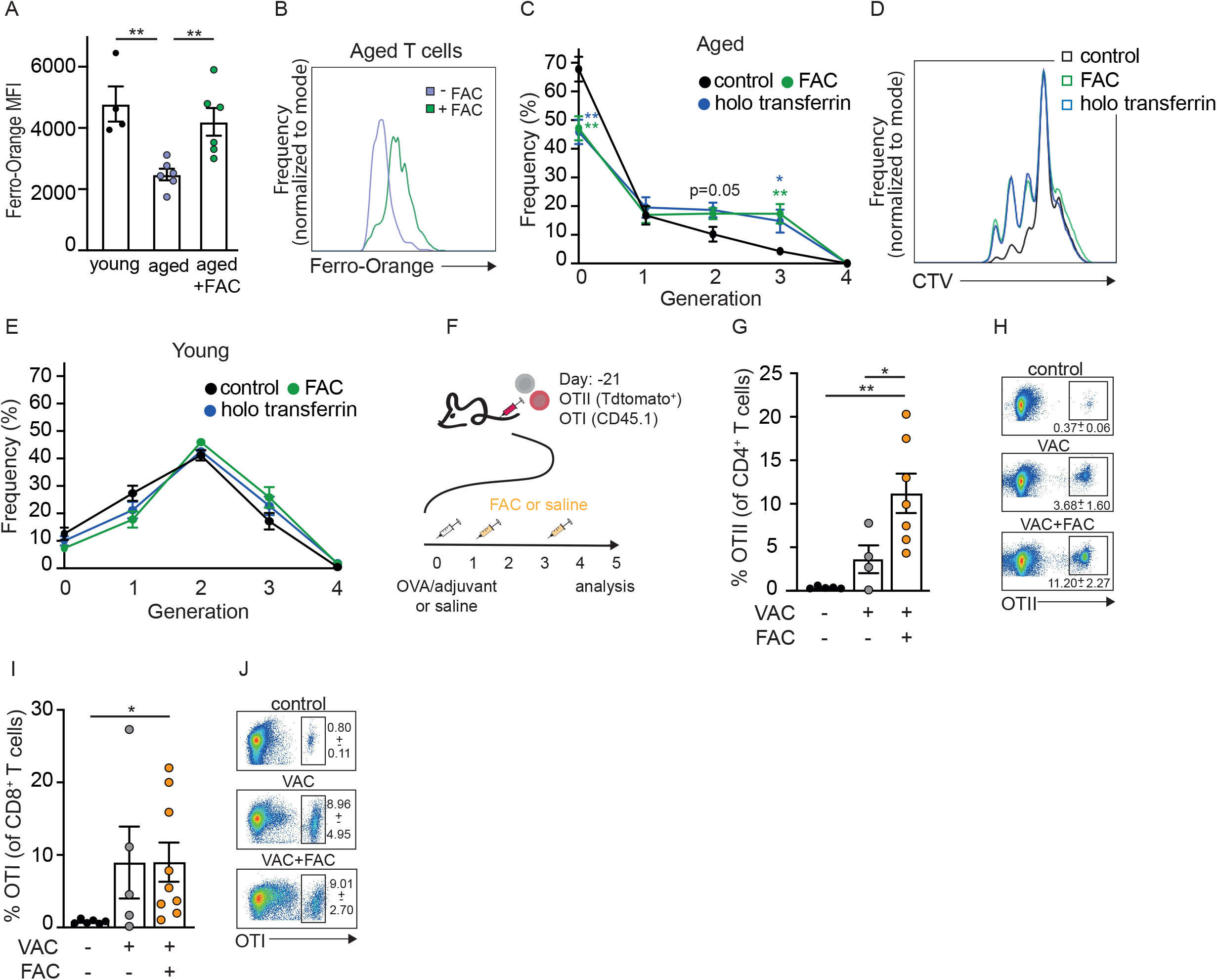
Iron supplementation rescued vaccination responses in aged T cells. (A) Ferro-Orange fluorescent intensity was quantified in young and aged T cells, and aged T cells supplemented with ferric ammonium citrate (FAC). (B) A representative plot shows intensity shift in labile iron content in aged T cells after supplementation with FAC. (C-E) young or aged T cells were loaded with CellTrace Violet and stimulated ex vivo with and without supplementation of FAC or holo transferrin, prior to analysis of proliferation. (C, D) analysis of aged T cells. (E) analysis of young T cells. (F) Schematic of experimental design. Aged C57Bl/7 wild type mice were inoculated with transgenic T cells bearing a known antigen specificity against ovalbumin (OVA). Each mouse was injected with a 1:1 mixture of OTII (TdTomato+CD4+) and OTI (CD45.1+CD8+) T cells. After two weeks, recipient mice were vaccinated with OVA/adjuvant intraperitoneally. Control mice received saline. On days 1 and 3 post-vaccination, vaccinated mice received i.v. infusions of FAC or saline. The mice were sacrificed on day 5 post-vaccination and T cell content in the spleen was analysed. (G) Quantitation and (H) a representative plot showing percentage of OTII+ T cells. (I) Quantitation and (J) a representative plot showing percentage of OTI+ T cells. Line graphs represent mean ± SEM. Data points (A,G,I) represent single mice . (*P<0.05, **P<0.01; unpaired student’s t-test).

The connection between iron availability and T cells response to vaccination was confirmed in a recent study using hepcidin-induce iron deficiency. T cells response to vaccination was diminished in mice treated with hepcidin and rescued by in vivo administration of FAC ^27^. We took a similar approach to test whether iron supplementation could improve T cells response to vaccination in aged mice. T cells with a known antigen specificity against ovalbumin (OVA) were isolated from transgenic young mice and injected into aged recipients. Each mouse received a mixture of OVA-specific OTI (CD8^+^CD45.1^+^) and OTII (CD4^+^TdTomato^+^) T cells. 3 weeks following cell transfer, recipient mice were vaccinated intraperitoneally with OVA emulsified in Alum adjuvant. Control mice were injected with saline. FAC was administered to vaccinated mice intravenously on days 1 and 3 following vaccination. Mice were sacrificed on day 5, and T cell content in the spleen was analysed by flow cytometry (Fig. 7F). Iron supplementation post vaccination, at times of high iron demand, improved proliferation of antigen specific CD4^+^ T cells (OTII; Fig. 7G, 7H). The proliferation of antigen specific CD8^+^ T cells was not significantly affected by FAC administration (Fig. 7I, 7J), in agreement with our data showing that these cells were less prone to iron deficiency (Figure 6).

## Discussion

Ageing manifests as a varied process among individuals and across distinct tissues within the same individual. Ageing in different tissues could differ in pace and propensity, and includes alterations in tissue structure and cellularity, changes in transcription profile, and proteome ^28–30^. Immune cells, including T lymphocytes are unique in the way that they are not limited to one tissue and instead are required to adapt to changes in their immediate microenvironment as they populate different tissues. Such adaptation is manifested by changes in the cells’ metabolic phenotype as T cells reshape their metabolism depending on the specific conditions and available resources in their host tissue. These include ad-hoc adaptations at a site of inflammation or within a solid tumour, and homeostatic adaptations of tissue resident lymphocytes to the specific conditions in their host tissue^31^. Our study shows how ageing processes in specific tissues shape the ageing trajectory of its resident T cells (Figure 1) and provide a new driving mechanism for known phenotypes of aged T cells, including reduced viability and low proliferative capacity. We found that exposure to the aged spleen microenvironment, enriched with senescent red blood cells and the products of their uncontained lysis, induces an adaptive response in T lymphocytes, manifested in an upregulation of proteins mediating the catabolism of haem and storage of excess iron (Figures 2,3). Ex vivo studies further highlight haem as a potent driver of T cell ageing phenotypes, as culturing young T cells in the presence of haem was sufficient to induce the in vivo observed phenotypes of elevated CD39 expression, reduced viability, and low proliferation (Figure 4). We further show that these all are purposeful adaptive changes that enable T cells to survive the hostile milieu of the aged spleen and resist ferroptosis, as aged T cells derived from spleens are significantly more resistant to treatment with ferroptosis inducers, such as RSL3 (Figure 5). Mechanistically, we show that T cells resist ferroptosis by restricting cellular labile iron pools (Figure 6). Iron is critical for DNA synthesis and iron deficiency is expected to inhibit proliferation, a known phenotype of age-related T cells dysfunction. Indeed, we show that addition of iron improves aged T cells proliferation in response to vaccination (Figure 7).

This study began with the isolation of pure naive T cells from young and aged mice. Given our interest in exploring age-driven changes in cellular metabolism, we sought to ensure comparability by utilizing known cell surface markers (CD25, CD62L, CD44) to purify naïve T cells. However, our findings demonstrate that cells collected from an aged spleen are not truly naïve as their proteome indicates cellular stress and inflammation. Many of the proteins overexpressed in aged ‘naïve’ T cells, remained significantly elevated in aged compared to young T cells, even after activation. These include proteins involved with protein maturation and folding, stress, inflammation, and T cells effector functions. Thus, our findings support inherent loss of proteostasis, in agreement with studies showing accumulation of misfolded proteins ^32,33^, reduced activity of proteolytic systems, like the proteasome ^34^ and autophagy ^35^, and altered chaperone expression ^36^ in aged T cells. Loss of proteostasis in spleen’s resident T cells is further supported by their increased cell size (^37^, Fig. 1). One potential mechanism could be mediated by ROS, as previous studies connect oxidative stress to alterations in proteostasis in aged cells ^38,39^, and we show that exposure to haem induces robust ROS production in T cells.

Interventions to rejuvenate T cell functions act on different modules of the proteostasis network. Those include inducers of autophagy and lysosomal activity, like mTOR inhibitors ^40,41^, Metformin ^42,43^, and spermidine ^35^, resulting in improved T cell responses and attenuated inflammation. Our study suggests that to some extent, these alterations in proteostasis could reflect adaptation to the aged microenvironment. Under such circumstances, rejuvenating these machineries could potentially expose the cells to toxic signals. For example, we found that developing resistance to ferroptosis is crucial for surviving in the aged spleen. We further show that T cells residing in the aged spleen are iron deficient despite high intracellular ferritin levels, suggestive of inability to release iron from its ferritin cage by ferritinophagy, a selective type of autophagy. We postulate that by avoiding ferritinophagy aged T cells avoid iron overload and ferroptosis, a mechanism previously described in senescent cells ^44^. Thus, combining therapies to improve proteostasis with interventions that reduce iron overload and haem accumulation in the aged spleen, could prevent lymphocytes loss by ferroptosis.

Haem is the reactive centre of multiple metal-based proteins. However, free haem has toxic properties like its ability to intercalate biological membranes, due to its hydrophobic nature, and the presence of the Fe-atom, which can drive ROS production through the Fenton reaction. Haem scavenging is mediated by different factors, including hemopexin, low-density lipoproteins, high-density lipoproteins, and serum albumin ^16^. Albumin is the most abundant protein in the serum. With ageing, albumin levels are dropping ^45,46^ alongside changes in its binding capacity ^47,48^. Administration of recombinant albumin prolonged lifespan and health span in mice ^48^. Our studies show that BSA mitigates the negative effects of haem and spleen extracellular components on T cells. In addition to its ability to sequester haem and its by-products^49^, albumin could protect cells from ferroptosis by increasing cellular cysteine levels ^50^. Another strategy for reducing haem and iron toxicity in the aged spleen is through a life-long iron restricted diet. As demonstrated by a recent study, reducing iron consumption preserved the function of red pulp macrophages, maintaining their ability to clear senescent erythrocytes and recycle iron through old age. In agreement, the spleens of iron-restricted aged mice had low amounts of haem, and iron depositions compared to their age-matched controls ^10^. Additional studies are required to test if and how this dietary intervention will affect spleen resident lymphocytes.

T cells residing in the aged spleen, and T cells acutely exposed to red blood cell lysate become resistant to ferroptosis. Our results suggest that this is mediated, in part, through restriction of labile iron pools, by reducing expression of transferrin receptor and upregulating ferritin; A mechanism supported by previous studies ^51,52^. However, other mechanisms could induce ferroptosis resistance in response to exposure to red blood cell lysate. Red blood cells are highly enriched in redox regulating enzymes (such as peroxiredoxin 2, glutathione peroxidase, catalase, and superoxide dismutase), and small molecule antioxidants (like α-tocopherol, glutathione, and ascorbic acid). It is possible that during an unregulated breakdown of red blood cells, some of these factors are released into the microenvironment, benefiting neighbouring cells by protecting against oxidative damage and ferroptosis ^53^. Red blood cells further contain high concentrations of IL-33 ^54^. When poured into the microenvironment, IL-33 could induce ferroptosis resistance through the ATF3/SLC7A11 axis ^55^. These compounds could be present in the interstitial-fluid enriched fraction of aged spleens, explaining its ability to promote ferroptosis resistance.

CD39 marks dysfunctional CD4^+^ T cells in human peripheral blood ^5^. We found that what induces CD39 overexpression on aged T cells is exposure to the milieu of the aged spleen. Moreover, while previous studies suggested that elevation of CD39 was mediated by the by- products of haem degradation: bilirubin ^56^ and CO ^57^, our use of an HO-1 inhibitor indicated that haem itself is a potent inducer of CD39 in aged T cells. We postulate that haem mediated CD39 expression is part of a regulatory mechanism that counteracts inflammatory signals in the aged spleen, and together with iron regulation, suppresses proliferation of aged T cells. Haemolytic conditions in the aged spleen pour ATP into the microenvironment ^20,58,59^. Extracellular ATP serves as a danger signal indicative of tissue damage, and induces proinflammatory signals, such as the inflammasome^19^. CD39, together with CD73 (both found to be elevated on T cells residing in the aged spleen) mediate the breakdown of ATP to adenosine which inhibits T cell proliferation ^21^.Similarly, this pathway plays a central role in mediating immunosuppression within the tumour microenvironment ^60^.

Our studies suggest a new approach for improving adaptive immunity in elderly, by targeting the hostile milieu of the aged spleen. We propose that reducing haem and iron toxicity in the aged spleen will alleviate T cell immunity and could improve immunosurveillance and everyday protection. We overcame the deleterious effect of the aged spleen by supplementing with iron, a relatively straight forward intervention in the context of vaccination, as we can anticipate the times in which iron is most needed. In opposed to chronic iron supplementation that could exacerbate iron accumulation in the tissue. Beyond ageing, these findings offer insight into potential mechanisms underlying other pathological scenarios involving haemolytic stress and accompanied by T cell dysfunction, such as sepsis ^61^, and sickle cell anaemia ^62^.

## Methods

### Mice

Young (7–10 weeks old) and aged (20-23 months old) C57BL/6JOlaHsd female mice were purchased from Envigo (Israel). For ageing experiments, retired breeders were purchased at 8 months and housed at the Technion animal facility for additional 12-15 months. Transgenic C57Bl/6 Rosa26^tdTomato/+^OTII mice were kindly received from Dr. Ziv Shulman (The Weizmann Institute of Science). C57Bl/6.SJLPtprcaPep3b;Ly5.1-Tg(TcraTcrb)1100Mjb/J (CD45.1 OTI) transgenic mice were kindly received from Dr. Michael Berger (The Hebrew University). All mice were housed in specific pathogen-free conditions at The Technion Pre–Clinical Research Authority and used in accordance with animal care guidelines from the Institutional Animal Care and Use Committee.

### T cell isolation and culture

T cells were harvested from mice spleens and lymph nodes (inguinal, axillary, brachial, mandibular, and jejunal), and purified by magnetic separation to obtain bulk CD3^+^- or CD3^+^CD4^+^ T cells or naïve (CD3^+^CD4^+^CD62L^+^CD44^lo^CD25^-^) T cells, using commercially available kits (StemCell). For isolating naïve T cells from aged mice, biotinylated anti-mouse/human CD44 (103004, biolegend), and biotinylated anti-mouse CD25 (130-049-701, Miltenyi Biotech) were added to the company’s premade antibody cocktail for negative selection of naive T cells, followed by a positive selection of CD62L^+^ naïve T cells on a magnetic column. Primary T cells were cultured at 37 °C and 5 % CO_2_ in RPMI media, supplemented with: 10 % FBS, 10mM Hepes, penicillin/streptomycin, and 0.035 % beta-mercaptoethanol. For activation, cells were cultured on plates pre-coated (overnight at 4°C) with anti-CD3 (2 μg/mL; BioXcell BE0001-1) and anti-CD28 (4 μg/mL; BioXcell BE0015-1). Resting cells were supplemented with 5 ng/ml recombinant murine IL-7 (Peprotech 217-17-10). In some experiments, T cell cultures were supplemented with hemin chloride, ammonium iron (III) citrate, BSA, or a spleen interstitial fluid-enriched fraction.

### Collection of spleen interstitial fluid-enriched fraction (referred to as SE)

Spleens from young or aged mice were harvested, and gently dissociated on a 70mM cell strainer, in 5mL PBS or RPMI, followed by a 5 min centrifugation at 1350 RPM, to separate cellular pellet from fluidic fraction, enriched with interstitial components.

### Red blood cell isolation and lysate

Whole blood was collected from mice via cardiac puncture into MiniCollect EDTA tubes (Greiner, 450531), diluted 1:1 in PBS, and subjected to leukoreduction using Hystopaque (Sigma, 10771) at 400g, without brake, for 30 minutes at room temperature. Red blood cell pellets were lysed by 4 freeze/thaw cycles.

### Flow cytometry

For cell-surface staining, T cells were suspended in a separation buffer (PBS containing 2 % FBS and 2 mM EDTA) and incubated for 20 min, on ice, with an antibody mix, supplemented with purified anti-mouse CD16/32. For intracellular staining, True-Nuclear™ Transcription Factor Buffer Set was used, following the manufacturer’s protocol. Staining for ferrous iron was done by incubation with 1µM of FerroOrange (dojindo) in HBSS at 37°C for 30 minutes in 5% CO2. Lipid peroxidation was assessed using BODIPY™ 581/591 C11 (Invitrogen), 5 µM in PBS at 37°C 5% CO2 for 30 minutes, or Liperfluo (dojindo) following the manufacturer instructions. Intracellular ROS was measured using by DCFDA / H2DCFDA - Cellular ROS Assay Kit (Abcam) according to the manufacturer’s instructions. All data were collected on the Attune NxT Flow Cytometer (Thermo Fisher) and analysed using FlowJo (BD).

A full list of flow cytometry reagents and antibodies:

| Antibody | Source | Catalog number |
| --- | --- | --- |
| Ferroportin/SLC40A1 | novus biologicals | 21502 |
| Alexa Fluor 647-AffiniPure Donkey Anti-Rabbit IgG (H+L) | Jackson | 711-605-152 |
| Alexa Fluor® 488 AffiniPure™ Goat Anti-Rabbit IgG (H+L) | Jackson | 111-545-144 |
| Alexa Fluor® 488 anti-mouse CD45.1 | Biolegend | 110717 |
| Alexa Fluor® 647 anti-mouse CD39 | Biolegend | 143807 |
| Alexa Fluor® 700 anti-mouse CD62L | Biolegend | 104426 |
| APC anti-mouse CD4 | BioLegend | 100412 |
| APC anti-mouse CD8a | BioLegend | 100712 |
| Brilliant Violet 421™ anti-mouse CD3 | BioLegend | 100227 |
| Brilliant Violet 421™ anti-mouse CD4 | BioLegend | 100438 |
| Brilliant Violet 421™ anti-mouse CD71 Antibody | Biolegend | 113813 |
| Brilliant Violet 421™ anti-mouse CD8a Antibody | Biolegend | 100737 |
| Brilliant Violet 510™ anti-mouse CD4 | Biolegend | 100449 |
| Brilliant Violet 510™ anti-mouse CD69 Antibody | Biolegend | 104531 |
| Brilliant Violet 510™ anti-mouse/human CD44 | Biolegend | 103043 |
| Brilliant Violet 605™ anti-mouse CD69 | BioLegend | 104529 |
| FITC anti-mouse CD25 Antibody | Biolegend | 102005 |
| FITC anti-mouse CD28 | Biolegend | 122007 |
| FITC anti-mouse CD3 | BioLegend | 100203 |
| FITC anti-mouse CD4 | BioLegend | 100406 |
| FITC anti-mouse CD73 | Biolegend | 127219 |
| FITC anti-mouse CD8a | BioLegend | 100706 |
| PE anti-mouse CD25 Antibody | Biolegend | 102007 |
| PE anti-mouse CD3 | BioLegend | 100206 |
| PE anti-mouse CD39 | BioLegend | 143804 |
| PE anti-mouse CD4 | BioLegend | 100407 |
| PE anti-mouse CD45.1 | Biolegend | 110707 |
| PE anti-mouse CD69 | BioLegend | 104508 |
| PerCP anti-mouse CD4 Antibody | Biolegend | 100537 |
| PerCP anti-mouse CD45 | Biolegend | 103129 |
| Purified anti-mouse CD16/32 | Biolegend | 101302 |
| Recombinant Rabbit Anti-Ferritin (FTH1) | Abcam | ab75973 |
| Recombinant Rabbit Anti-Haem Oxygenase 1(HO-1) | Abcam | ab52947 |

### Untargeted, whole-cell Proteomics

Naïve T cells (CD4^+^CD62L^hi^CD44^lo^CD25^-^) were purified from young and aged mice spleens. Cell pellets were frozen immediately or after 24 hr stimulation, followed by protein extraction and digestion. 2 μg protein per sample were analysed by LC-MS/MS on Q-Exactive plus (ThermoFisher). Collected data were processed using Maxquant (1.6.17.0; Mathias Mann, Max Planck Institute) and identified against the mouse proteome (Uniprot database (Jan 2020)), and a decoy database. Statistical analysis was performed using Perseus (1.6.14.0). Pathway enrichment analysis was performed using GSEA (Broad Institute and UC Sn Diego).

### Adoptive T cell transfer

Young T cells were isolated from spleens of young C57Bl/6 Rosa26^tdTomato/+^ or C57Bl/6 Rosa26^tdTomato/+^OTII or CD45.1 OTI transgenic mice (8-12 weeks-old) by magnetic isolation (StemCell). 5X10^6^ cells were transferred i.v., into wild type C57Bl/6 young or aged recipients.

### Histology

Spleens harvested from young and aged mice were fixed with 4 % PFA, processed in Tissue Processor (Leica TP1020, Germany), paraffin embedded, and sectioned (4μm; Leica RM2265 Rotary Microtome, Germany). Sections were stretched on a warm 37 °C water bath, collected into slides, and dried at 37 °C overnight. Sections were processed for H&E and Prussian blue staining. Slides were scanned by 3DHistech Panoramic 250 Flash III and visualized using the CaseViewer software (3DHISTECH).

### Immunohistochemistry

Tissue dissection, fixation, embedding, and sectioning was performed as previously described^63^, and stained with AF647 anti-CD169 (Biolegend). Imageing was performed using an LSM710 AxioObserver microscope.

### Quantitative, real-time PCR (qPCR)

Total RNA was extracted and from CD3^+^ T cells, using QuickRNA Micro prep Kit (ZYMO RESEARCH). cDNA was synthesized using the High-Capacity cDNA RT kit (Applied Biosystem). Quantitative PCR was run on QuantStudio™ 3 Real-Time PCR System, 96-well (Applied Biosystem), using Fast SYBR Green Master Mix (Applied Biosystems).

Primers used: *Blvra,* F-AAGATCCCGAACCTCTCTCT, R: TTATCAAGGCTCCCAAGTTCTC; *Blvrb,* F: AAGCTGTCATCGTGCTACTG, R: CAGTTAGTGGTTGGTCTCCTATG; *Fth1*, F: CGTGGATCTGTGTCTTGCTTCA, R: GCGAAGAGACGGTGCAGACT; *Hmox1*, F: GTTCAAACAGCTCTATCGTGC, R: TCTTTGTGTTCCTCTGTCAGC; *Rps18*, F: CCGCCATGTCTCTAGTGATCC, R: GGTGAGGTCGATGTCTGCTT,

### Vaccination and in vivo iron supplementation

Young T cells were isolated from Rosa26^tdTomato/+^OTII or CD45.1 OTI transgenic mice by magnetic separation, and equally mixed at a 1:1 ration. A total of 4M cells were inoculated by tail vein injection into wild-type C57Bl/6 young or aged recipients. 3 weeks follow cell transfer, recipient mice were injected intraperitoneally (i.p.) with OVA albumin (vac-stova; InvivoGen) adsorbed in 40% Alum adjuvant (Alu-Gel-S; SERVA) or with 0.9% saline. On days 1 and 4 following vaccination, some mice received injections of ferric ammonium citrate (Ammonium iron(III) citrate, brown (Thermo Fisher); 900μg/mouse). Mice were sacrificed on day 5 following vaccination, the spleens were harvested and analysed by flow cytometry to quantify OTII tdTomato^+^ and OTI CD45.1^+^ T cells.

### Statistical analysis

Was Performed using the Prism software.

## Acknowledgements

The authors thank Dr. Tamar Ziv from The Smoler Proteomics Centre. Ifat Gavish-Abramovich and Dr. Bella Agranovich from The Perlmutter Metabolomics Centre. Prof. Michael Berger, and Prof. Ziv Shulman for providing mouse models. Prof. Esther Meyron Holtz, Prof. Katarzyna Mleczko-Sanecka and Prof. Michael Berger for discussions. Ms. Viktoria Zlobin and Dr. Amit Avrahami from the Technion Preclinical Authority for maintaining aged mouse colony and help with in vivo studies. The research was Funded by the European Union, by an ERC-Stg grant awarded to NRH. Views and opinions expressed are however those of the author(s) only and do not necessarily reflect those of the European Union or European Research Council.

## Author contributions

Conceptualization, N.R.-H. and D.E.; Methodology and investigation, N.R.-H., D.E., H.O., L.W., O.A.; Writing, N.R.-H. and D.E.; Visualization, N.R.-H. and D.E.; Funding Acquisition, N.R.-H.

## Declaration of interests

The authors declare no competing interests.

## Notes

### Competing Interest Statement

The authors have declared no competing interest.

