## Extended data figures for "Haem toxicity in the aged spleen impairs T-cell immunity through iron deprivation"

Extended data figure 1

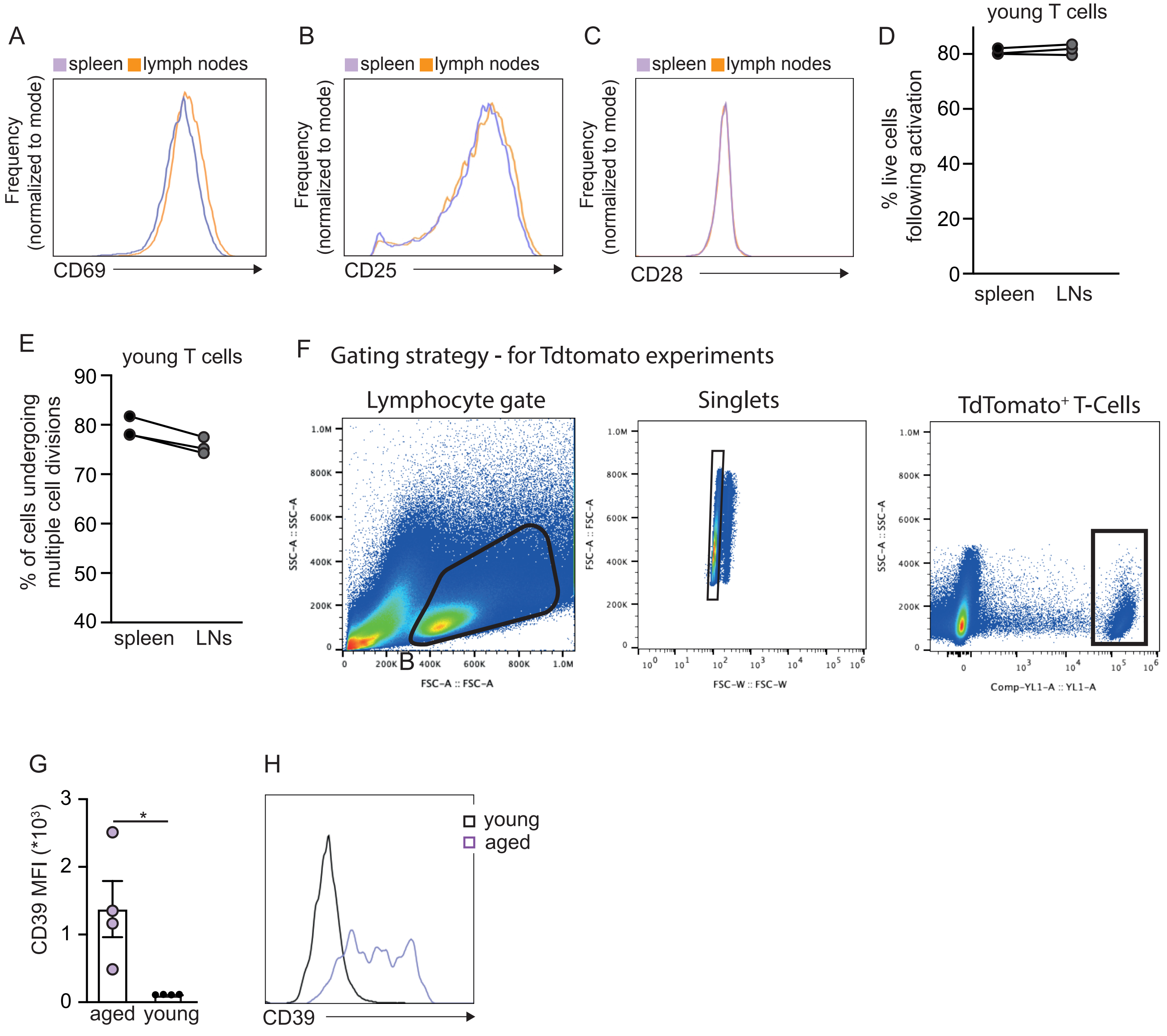


**Extended data figure 1:**
Naïve T cells (CD4+CD25-CD62L+CD44lo) were purified from the spleen or peripheral lymph nodes (LNs) of aged mice (20-22 months old). The cells were kept in two separate pools and stimulated ex vivo for 48 hr using plate bound anti-CD3/anti-CD28, prior to analysis by flow cytometry to assess expression of (A) CD69), (B) CD25 and (C) CD28. A similar experiment compared (D) viability and (E) proliferation between T cells isolated from the spleens and LNs of young (8 weeks old) mice. (F) Gating strategy for adoptive transfer experiments described in Figure 1G. (G) Quantitation and (H) a representative plot showing CD39 expression on CD4+TdTomato+ T cells derived from the spleens of young and aged mice, following ex vivo activation. Each data point represents data collected from an individual mouse. Bar graphs show mean ± SEM. (*P<0.05, paired student's t test when comparing cells derived from spleen and LNs of the same mouse, and unpaired student's t test when comparing values across age groups).

Extended data Figure 2

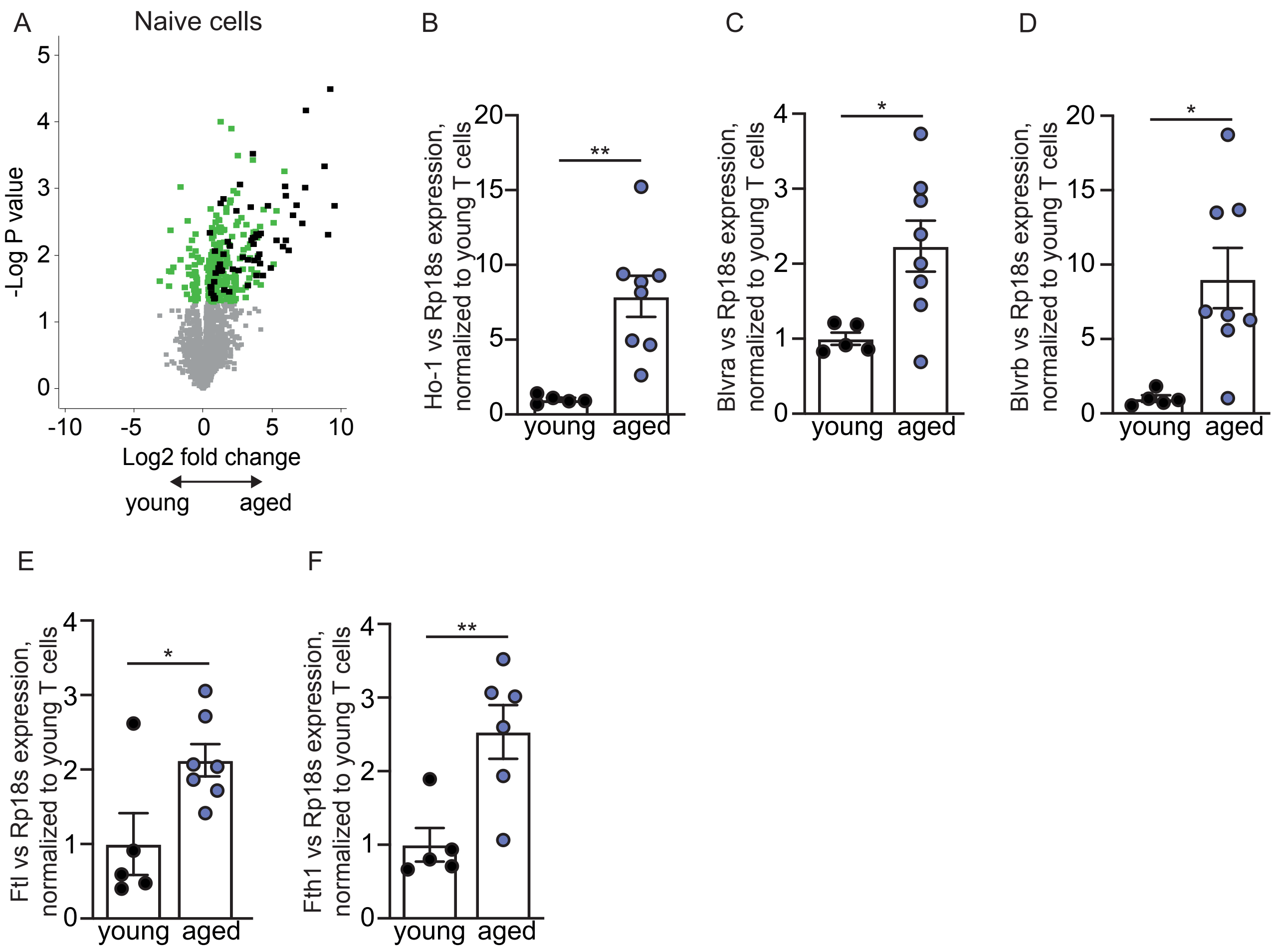


**Extended data figure 2:**
(A) A volcano plot showing differences in protein levels between naive young and aged T cells. Green signifies statistical significance. Black highlights proteins that remained significantly higher in activated aged compared to young T cells. (B-F) CD3+ T cells were isolated from the spleens of young and aged mice, processed, and analyzed by real-time qPCR to measure expression of genes encoding proteins involved in heme detoxification: (B) Ho1, (C) Blvra, (D) Blvrb, (E) Ftl, (F) Fth1. Each data point represents data collected from a single mouse. Bar graphs show mean ± SEM. (*P<0.05, **P<0.01; unpaired student's t test).

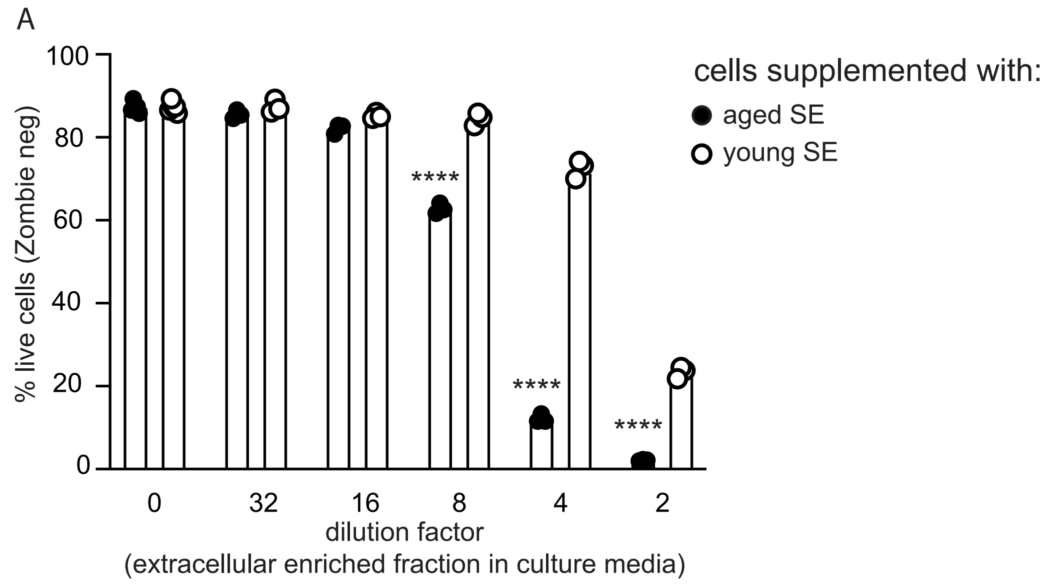


**Figure S3:**
(A) Young T cells derived from 3 mice were pooled together and cultured ex vivo in media supplemented with spleen's interstitial-fluid enriched fraction (SE) from young or aged spleens, in increasing concentrations. Cell survival was analyzed using the Zombie Aqua reagent. Data point represent technical repeats. Bar graphs show mean ± SEM. (****<0.0001; unpaired student's t test, the two conditions in each dilution).

### Extended data Figure 4

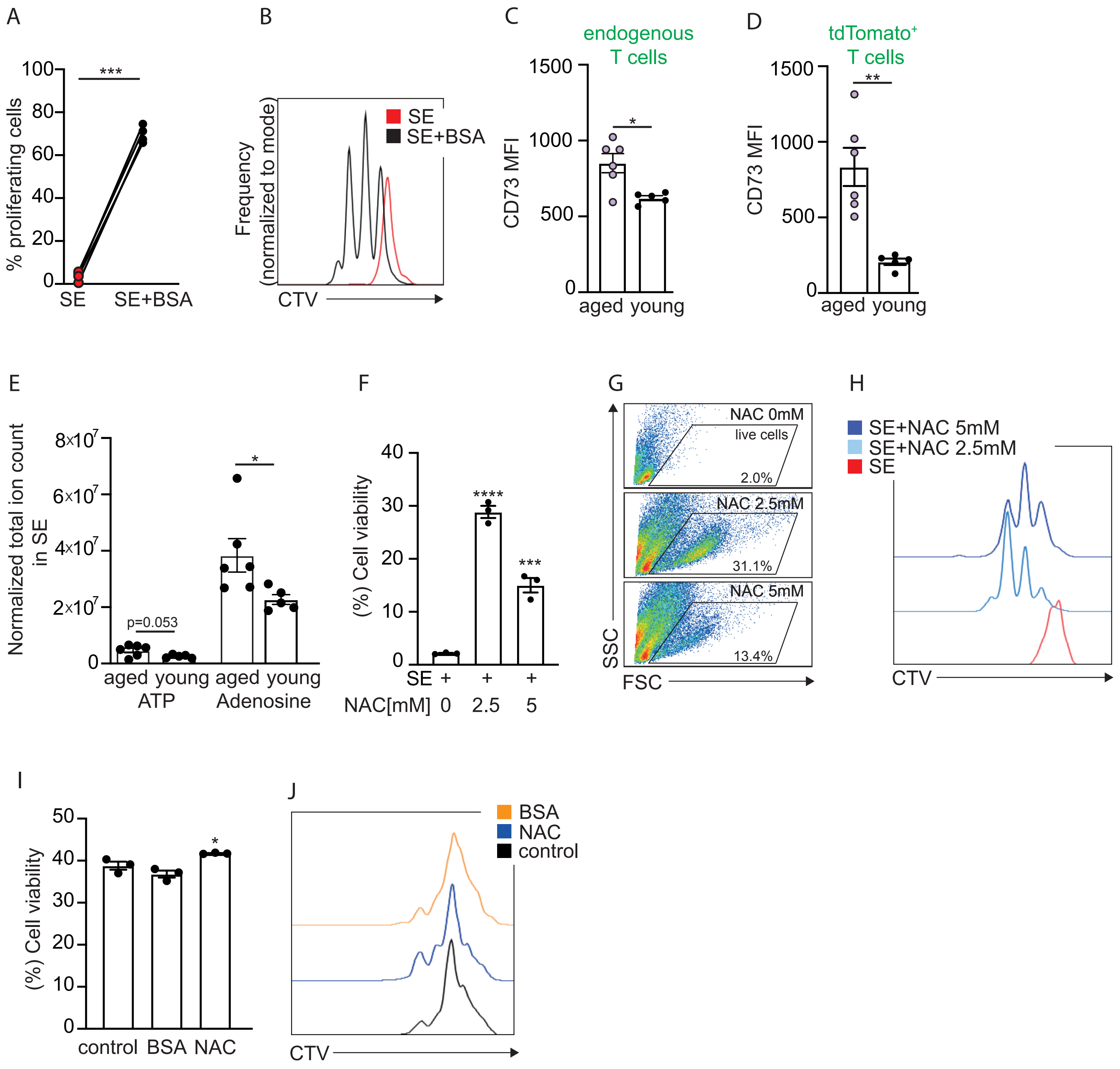


**Extended data figure 4:**
young T cells were loaded with CellTrace Violet (CTV), and stimulated ex vivo using plate-bound anti-CD3/anti-CD28, in media supplemented with aged spleen's interstitial-fluid enriched fraction (SE) in a 1:4 dilution. (A,B) The effect of BSA supplementation on proliferation was assessed by flow cytometry. (C,D) CD73 MFI on endogeneous (TdTomato-; C) and transferred (TdTomato+; D) T cells derived from the spleens of young and aged recipient mice. See Fig. 1G for experimental design. (E) Quantitation of ATP and adenosine in SE from young and aged mice. Showing normalized total ion counts. (F-H) Young CD3+ T cells were treated with SE in the presence of N-acetyl cysteine (NAC). Cell viability (F, G) and proliferation (H) were assessed by flow cytometry.
Aged T cells were stimulated ex vivo in the presence of BSA or NAC. (I) Cell viability and (J) proliferation were analyzed by flow cytometry. Each data point represents data collected from a single mouse. Bar graphs show mean ± SEM. (*P<0.05, ***P<0.001, ****<0.0001; unpaired student's t test: comparing each condition to control cells; paired student's t-test (A) comparing each sample to its untreated control).

Extended data Figure 5

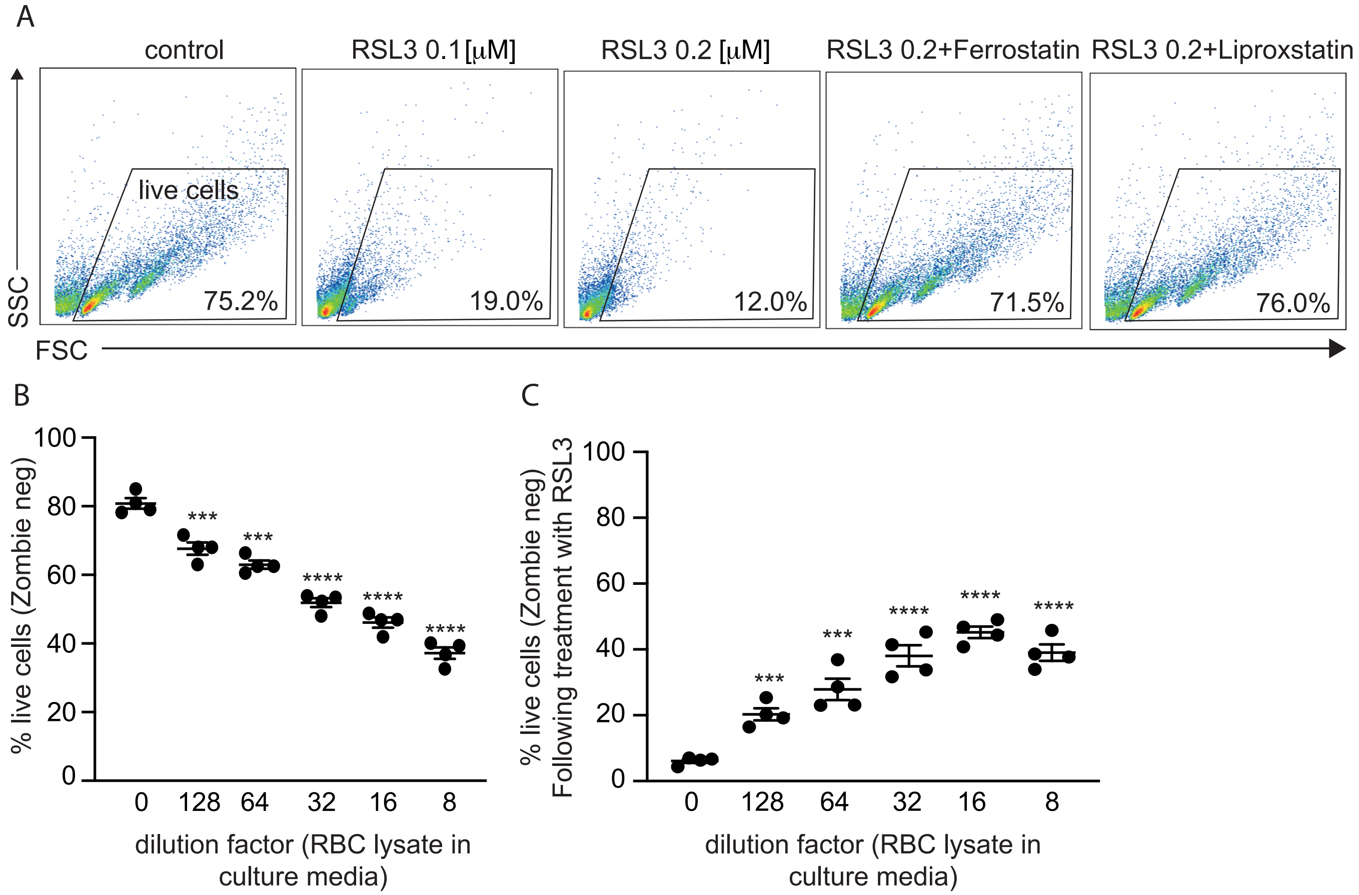


**Extended data figure 5:**
(A) Young T cells were cultured in media containing the indicated concentrations of the ferroptosis inducer, RSL3 ± Ferrostatin or Liproxstatin, two commercially available inhibitors of ferroptosis. Representative FACS plots with “live cells” gate are presented. (B) Young T cells were cultured in media containing red blood cells (RBC) lysate in the indicated concentrations. Cell viability was determined using the Zombie reagent and flow cytometry. (C) A similar experiment as in (B), with the addition of RSL3 to growth media. Each data point represents data collected from a single mouse. Bar graphs show mean ± SEM. (***P<0.001, ****<0.0001; unpaired student’s t test: comparing each condition to untreated control cells).

Extended data Figure 6

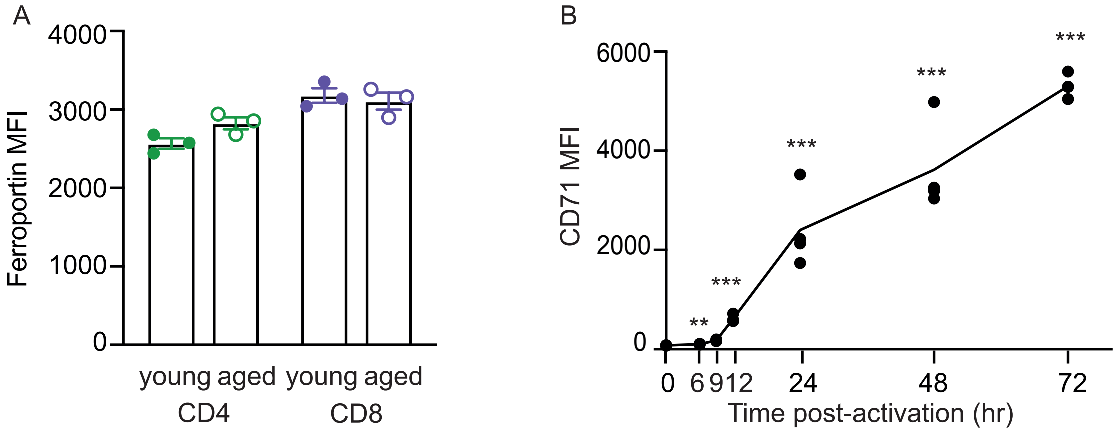


**Extended data figure 6:**
(A) Ferroportin expression levels in young vs. aged CD4+ and CD8+ T cells. Analyzed by flow cytometry. (B) Young T cells were stimulated ex vivo using plate bound anti-CD3/anti-CD28. CD71 expression was measured by flow cytometry at the indicated times post-activation. Data points indicate single mice. (**P<0.01, ***<0.001; unpaired student's t test: comparing each condition to control cells).

Supplementary Figure 7

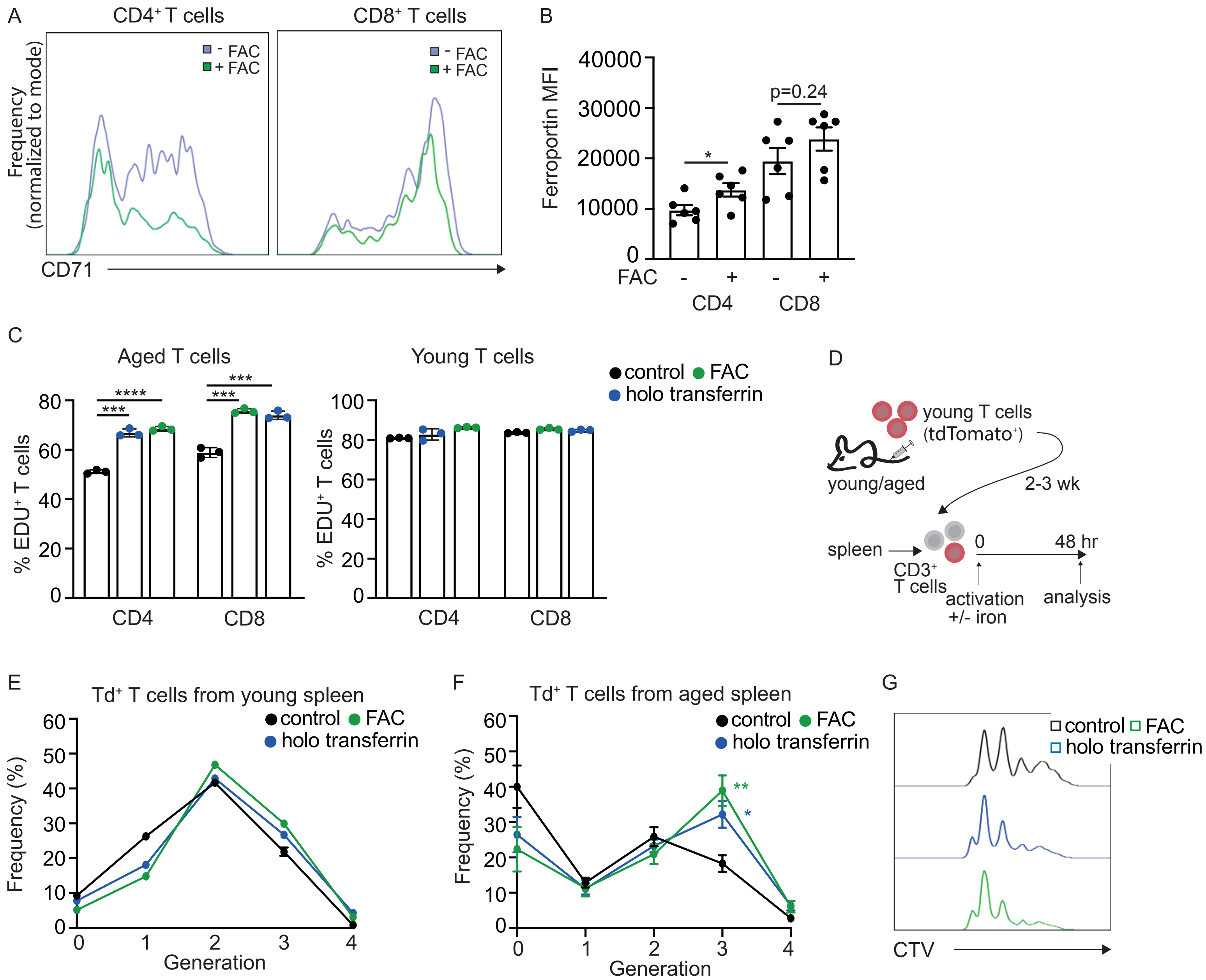


**Extended data Figure 7:**
(A) representative FACS plots showing CD71 expression on CD4+ and CD8+ T cells with and without supplementation of ferric ammonium citrate (FAC) to culture media.(B) Ferroportin levels in T cells activated with and without supplementation with ferric ammonium citrate (FAC). (C) Analysis of EDU (5-ethynyl-2'-deoxyuridine) incorporation into T cells activated with and without iron supplementation. (D) schematic of experimental design. Young TdTomato+ T cells were transfused into aged or young recipients. 2-3 weeks following the transfer, the mice were sacrificed, and T cells were loaded with CellTrace Violer, and stimulated ex vivo with and without iron supplementation: FAC or holo transferrin. (E) analysis of proliferation of TdTomato+ T cells isolated from young mice. (F) analysis and (G) representative plot depicting proliferation of TdTomato+ T cells isolated from aged mice. (*P<0.05, **P<0.01, ***P<0.001, ****P<0.0001; unpaired student's t-test comparing each condition to control cells).
